# Chemical Inhibition of Splicing-Related Protein Kinases Reveals Phosphorylation-Driven Regulation of RNA Alternative Splicing in *Arabidopsis* Seedlings

**DOI:** 10.1101/2025.03.24.644839

**Authors:** Maria Camila Rodriguez Gallo, Cameron Murray, Curtis Kennedy, Mohana Talasila, Devansh Bhatt, Joselyn Steyer, Mark Glover, Glen R. Uhrig

**Affiliations:** Department of Biological Sciences, University of Alberta, Edmonton, Alberta, Canada; Department of Biochemistry, University of Alberta, Edmonton, Alberta, Canada; Department of Computer Science, University of Alberta, Edmonton, Alberta, Canada

**Author notes:** **Corresponding Author:** Richard Glen Uhrig, Department of Biological Sciences, CCIS Building rm 5-114 / 5-106, University of Alberta, Edmonton, Alberta, Canada, T6G 2E9.

**Keywords:** *Arabidopsis thaliana*, Chemical Inhibitors, Alternative Splicing, Serine/Arginine Protein Kinases, Multi-Omics, Root Growth and Development

## Abstract

Serine / arginine (SR) protein kinases (SRPKs) are capable of transmitting external signals to the spliceosome by phosphorylating SR proteins. In plants, few studies have looked at the regulation of RNA splicing by post-translational modification (PTMs), despite the spliceosome and splicing factor proteins exhibiting extensive phosphorylation. Many of these PTM events have the potential to dramatically change the expression of various genes, such as those involved in abiotic stress. Here, we sought to explore the regulatory function of *Arabidopsis thaliana* SRPKs in the context of RNA alternative splicing. To do so, we utilized four well-known and well-characterized chemical inhibitors specifically designed to target and inhibit SRPK enzymatic activity. Our data found that SRPK chemical inhibitors SPHINX31 and SRPIN340 induced shorter root development phenotypes and an abolishment of root hair formation. Using a multi-omics approach combining transcriptomics and phosphoproteomics, we find extensive changes in the splicing of genes involved in root development, RNA splicing, cytoskeletal organization, and cell differentiation. We also reveal splicing factors exhibiting differential alternative splicing as well as a down-regulated in their phosphorylation status. Overall, our data indicate that AtSRPKs phosphorylate diverse splicing factors and influence the alternative splicing of genes involved in root development and a wide-range of related pathways.

## INTRODUCTION

Alternative splicing (AS) is a crucial co- and post-transcriptional mechanism that adds an additional regulatory layer to gene expression. AS has recently gained significant attention due to its role in enhancing genetic diversity and adaptability in response to environmental stresses. In humans, exon skipping (ES) is the most prevalent AS event and is thought to diversify the proteome by producing multiple mRNA isoforms from a single gene locus (1, 2). Beyond diversifying the proteome, another major function of AS is to regulate transcript and protein abundance though two major mechanisms: nuclear retention of alternative transcripts (3–5) and mRNA degradation via the nonsense mediated decay (NMD) pathway (6, 7). Unlike humans, plants favour intron retention (IR) (∼60% of all AS are IR events), an approach that has been hypothesized to modulate transcript levels and thus protein abundance in response to environmental stress (8, 9). The primary purpose of global IR has been commonly thought to trigger NMD pathway through the introduction of premature stop codons (PTCs) in mRNA transcripts (10–13). Recent studies have found that IR transcripts are predominantly retained in the nucleus and then spliced when the transcript is required at a later time, essentially utilizing IR as a spatial-temporal form of gene regulation (3). In particular, a 2024 study, found IR to be a mechanism for segregating photomorphogenesis-related mRNA transcripts in the nucleus in response to light (3). Additionally, in human cells, phase separation condensates can also act as a spatial-temporal regulation of AS splicing (14), wherein certain conditions induce the formation of splicing factor nuclear condensates affecting selection of specific splice sites (15), RNA-binding (16), and splicing efficiency (17). While condensates in the context of AS are relatively understudied in plants, emerging research shows that phase separation of specific splicing factors controls RNA splicing, which in turn affects growth and development (18).

In plants, evidence linking AS to environmental stress has largely been gathered through the study of specific mRNA transcripts and pathways that induce AS (19–21). Researchers have recognized that understanding AS regulation is essential for uncovering how plant cells modulate gene expression in response to various environmental signals, however, the functional outcomes of these splicing events ultimately depend on the proteins produced. Yet, few studies have investigated the relationship between the AS transcripts and its effects on downstream protein-level changes, with even fewer having examined the higher-order regulatory components that drive these AS events, such as the post-translation modification (PTM) events activating splicing proteins.

The spliceosome, a large multi-protein complex, is predominantly regulated through PTMs, such as protein phosphorylation (22–24). For example, U1 small nuclear ribonucleoprotein (snRNP), necessary for the formation of the early spliceosome and initiator of splicing events, is the most highly phosphorylated Arabidopsis snRNP (22). Furthermore, phosphorylation of splicing factors, such as serine arginine (SR) proteins, and heterogenous RNPs (hnRNPs) are highly phosphorylated and are known to induce and repress constitutive and AS (25–27). The high level of spliceosome phosphorylation indicates the presence of major regulatory networks between protein kinases transducing environmental signals to their spliceosome substrates. SR proteins are highly phosphorylated on C-terminal serine and arginine-rich domain by splicing-related protein kinases, allowing them to bind to regulatory elements and act agonistically on AS (28–30). In general, inhibiting splicing-related protein kinases would prevent the phosphorylation of serine / arginine (SR) proteins, leading to aberrant RNA splicing. Since the full regulatory role of SR proteins in splicing pathways remains unresolved, the specific mechanisms by which these splicing-related protein kinases regulate individual transcripts, as opposed to their broader effects on AS, remains to be studied.

Human SR protein kinases (HsSRPKs) are activators of SR proteins, thus changes to HsSRPKs phosphorylation capabilities lead to modifications of AS patterns related to DNA repair, cell cycle control, and apoptosis (31). Further, HsSRPKs and CDC-like kinases (HsCLKs) have been implicated in human angiogenic diseases (32–35). For example, HsSRPKs regulate the splicing of a pro-angiogenic gene (VEGF-A165) through phosphorylation of serine/arginine splicing factor 1 (SRSF1), inducing the translocation of SRSF1 from the cytoplasm to the nucleus and consequently binding to the VEGF-A pre-mRNA splice site, enabling the AS of the pro-angiogenic isoform (36). As such, the usage of chemical inhibitors to target HsSRPKs and HsCLKs have be extensively studied and have been commonly used in therapeutic settings, some of which have become FDA approved anti-cancer therapies.

Chemically inhibiting SRPKs allows for the broad study of induced dysregulation of phosphorylation-mediated splicing over the short term (days), as opposed to a constitutive abolition of only one splicing kinase gene through the use of single gene loss-of-function plant lines, where potential genetic redundancy may mask the scope of their involvement in plant cell regulation (37). Although chemical inhibitors are a common tool in studying aberrant AS in human cells, they are notably underutilized in plant systems. Only recently have plant researchers taken advantage of chemical inhibitors originally designed for human cells. For example, a handful of studies have used pladienolide B (PB), herboxidiene (GEX1A), and spliceostatin C (SPS) as splicing inhibitors in plants and showed their utility in studying post-transcriptional regulation of stress responses to uncover the interplay between AS and abiotic stress (38–40). Interestingly, these chemical inhibitors have the potential to serve as new herbicides and to be used concurrently with engineered plants resistant to these splicing inhibitors (40). In particular, GEX1A has been proposed as a herbicidal agent against various weed species, while also being inert to wheat (41)while in rice, SPLICING FACTOR 3B1 (SF3B1) has been engineered to resist inhibition through CRISPR-Cas9 based editing (42). Here, we tested four different chemical inhibitors specifically developed to have high potency and specificity towards HsCLKs and HsSRPKs: SPHINX31, Leucettine L41, Alectinib, and SRPIN340. While chemical inhibitors targeting the core spliceosome have been used in plant research, to our knowledge, splicing protein kinase inhibitors have not yet been employed in plants, highlighting a promising opportunity to explore the global regulation of AS at the PTM level.

Previously, we found three highly conserved families of splicing-related protein kinases, five SERINE ARGININE PROTEIN KINASES (SRPK1-5), three ARABIDOPSIS FUS3 COMPLEMENT (AFC1-3), and three PRE-mRNA PROCESSING FACTOR 4 (PRP4Ka-c) that are highly conserved across eukaryotes We therefore first determined if these splicing-related protein kinases are conserved relative to their human orthologs at both the sequence and structural level. Additionally, we examined whether the amino acids required for binding the chemical inhibitors are also present. When applied to *Arabidopsis thaliana* seedlings, these inhibitors revealed significant root growth phenotypes, with the most dramatic reduction in root growth at the lowest applied SPHINX31 concentration, which also resulted in a near abolishment of root hair formation. Subsequent use of transcriptomic and phosphoproteomic analysis then uncovered molecular events driving the root phenotypes, with these analyses revealing differential AS (DAS) of transcripts related to root development, splicing factors, and cell differentiation along with a down-regulation of phosphosites on proteins involved in RNA splicing, cell-cycle, meristem differentiation and root cell elongation, amongst others. Together, these data indicate that the splicing-related protein kinases, particularly AtSRPKs, are not only highly conserved in their sequence to the human orthologs, but also functionally conserved, suggesting plant SRPKs may have ancestral function in eukaryotic cells. Further, our data involving AtSRPKs suggests that SRPKs in plants, similar to HsSRPKs, regulate a substantial number of biological processes beyond their canonical role as regulators of SR proteins.

## MATERIAL AND METHODS

### Amino Acid Similarity Across Taxa

For amino acid similarity, all peptide sequences of human SRPKs (HsSRPK1: Q96SB4, HsSRPK2: P78362, HsSRPK3: Q9UPE1), Arabidopsis SRPKs (AtSRPK1: AT4G35500, AtSRPK2: AT2G17530, AtSRPK3: AT5G22840, AtSRPK4: AT3G53030, AtSRPK5: AT3G44850), and select photosynthetic organisms were aligned using MAFFT version 7 (https://mafft.cbrc.jp/alignment/server/). Percent conservation of each amino acid was generated and visualized using JalView vr. 2.11.3.0 (https://www.jalview.org).

### Percent Similarity of HsSRPKs and Arabidopsis Protein Sequences

Human and Arabidopsis proteins compiled amino acid sequences were aligned using MAFFT LINS-I with default settings. Resulting alignment was inputted through Sequence Identities and Similarities (SIAS) tool (http://imed.med.ucm.es/Tools/sias.html) with BLOSUM 62 as the scoring matrix. The length of the multiple sequence alignment (MSA) was used as the sequence length denominator used in the percent identity equation.

### Multiple Sequence Alignment

HsSRPK1 peptide sequence was searched against the Arabidopsis proteome using JGI Phytozome pBLAST tool (https://phytozome-next.jgi.doe.gov/blast-search) with default settings. Negating AtSRPKs, AtAFCs, and AtPRP4Ks hits, we selected the top candidates for MSA, which included: AtDYRK-2a (AT1G73460), AtDYRK-2b (AT1G73450), AtDYRKP-1 (AT3G17750), DYRKP-3 (AT2G40120) and AtYAK1 (AT5G35980). In addition, the SnRK2.2 (AT3G50500) peptide sequence was included in the multiple sequence alignment as an Arabidopsis protein kinase whose kinase activity has been inhibited by chemical inhibitors unrelated to those used here (44). R packages msa, Biostrings, and BiocManager were used to generate and visualize the MSA with ClustalOmega method.

### Structural Alignment of Human SRPK to Arabidopsis Protein kinases

Tertiary structures of AtSRPK1 (F4JN22), AtSRPK2 (Q9SHL5), AtSRPK3 (Q9FE52), AtSRPK4 (F4J982), AtSRPK5 (Q9FYC5), AtAFC3 (P51568), AtDYRK-2A (F4HQ88), AtSnRK2.2 (Q39192), YAK1 (Q8RWH3), were sourced from AlphaFold prediction entries (https://alphafold.ebi.ac.uk). Structures with predicted local distance difference test (pLDDT) scores less than 70 were removed for structural alignment to HsSRPK1 crystal structure (PDB: 1WAK) in PyMol vers. 3.1 (https://www.pymol.org). A root mean square deviation (RMSD) score was provided on aligned atoms by PyMol. Percent similarity across the entire peptide sequence between HsSRPK1 and the Arabidopsis protein kinases were generated by performing a pairwise protein alignment using the EMBL EMBOSS needle global pairwise alignment tool (https://www.ebi.ac.uk/jdispatcher/psa/emboss_needle). TM-score was generated by using TM-align (https://zhanggroup.org/TM-align/) with default settings.

### Docking Calculations of Chemical Inhibitors on Arabidopsis Proteins

The tertiary structures of AtSRPK1 and AtSRPK4 as representative Group I and Group II SRPKs (43), were sourced from the AlphaFold predictions listed under UniProt entries F4JN22 and F4J982 as their structures were not available in the Protein Data Bank (PDB). Molecular docking simulations employed the amber94 force field (45) as implemented in AutoDock Vina (46). Subsequent analyses of docking results were performed using LigPlot+ v1.4 (47).

### Splicing Inhibitor Root Length and Root Hair Phenotypic Assay

Sterile Arabidopsis wild-type Col-0 (WT) were imbibed on 0.5 X MS (Murashige & Skoog - Caisson Laboratories Inc.; MSP01) + 0.8% plant agar (Caisson laboratories Inc. Phytoblend™; PTP01) media and placed at 4°C for 3 days. Seedlings were grown under a 12h : 12h light : dark photoperiod consisting of 100 µmol/m²/s light from fluorescent bulbs and 22°C in custom-build black vertical plate boxes to shield roots from light. After 4 days, seedlings were transferred to 0.5 X MS - agar plates containing inhibitor media (SPHINX31; Cayman chemical; 1818389-84-2, SPRIN340; Cayman Chemical; 218156-96-8, Leucettine L41; Cayman Chemical; 1112978-84-3, Alectinib; Cayman Chemical; 1256580-46-7) or control media containing DMSO (0.1 % v/v). Seedlings were grown on treatment plates for 7 days with root length measured daily. A one-way ANOVA was performed to determine significant root growth differences between experimental and control conditions.

For root hair analysis, the seedlings were imaged under a dissecting microscope (Leica L2) at 10X magnification. Images were taken using a digital camera (Ningbo Icoe Commodity Co.; HDCE-90D) attached to the microscope. The imaged area is located approximately 2 cm above the root tip which correlates to the zone of maturation (Supplemental figure 1). Root hairs were quantified using automatic counting with ImageJ. Images were processed as follow: convert image to 8-bit > adjust threshold (default, B&W, select dark background, 255 and 255) > analyze particles (150-infinity-1.00, outlines).

### Plant Growth for Transcriptomic and Proteomic Analysis

Arabidopsis seed were sterilized and then stratified at 4°C for 4 d in the dark. Seeds were then plated on germination media 0.5 X MS + 0.8% and left to germinate for 10 d as described above. Seedlings were then transplanted to either control plates (DMSO) or inhibitor plates (8 µM SPHINX31). Again, in custom-build black vertical plate boxes to shield roots from light. Seedlings were harvested after 2 d and 7 d of treatment at ZT23 (before dawn). Seedlings were then harvested into tubes and flash frozen in liquid nitrogen and stored at −80°C prior to extraction. Seedlings were ground using a GenoGrinder (SPEX Sample Prep) at 1200 rpm for 30 sec, with 100 mg of the 2 d tissue aliquoted for RNA analysis and 400 mg aliquoted for proteomic analysis.

### RNA Extraction, cDNA Synthesis, and RNA Sequencing

Total mRNA was extracted from 100 mg of ground tissue. RNA was extracted using Direct-zol-96 MagBead RNA kit (Zymo research; R2100). Degraded RNA was checked by using a Tapestation 4150 (Agilent) following manufacturer’s protocol. RNA-seq library was prepared by applying Smart-seq2 single-cell method on bulk RNA. cDNA was created by following the protocol previously described (48) with the following adjustments: 1 µL of lysate was transferred from lysis plate into a new plate containing the RT mix. Preamplification of cDNA used 23 PCR cycles and was purified using Agencourt Ampure XP beads (Beckman Coulter; A63880) with a modified bead: DNA ratio of 0.8 X. The quality of cDNA was checked using a NGS Fragment High Sensitivity Analysis Kit (Advanced Analytical) and a Fragment Analyzer (Advanced Analytical). The cDNA concentration was measured using a qubit High sensitivity dsDNA Kit. Libraries were prepared using a Nextera XT DNA Library Preparation Kit (Illumina), using a standard protocol but with all reaction volumes reduced by 1/10 to accommodate the automation of the prep on the Echo LabCyte liquid handler (Beckman Coulter). Libraries were purified using Agencourt Ampure XP beads (Beckman Coulter). Size distribution of library pools was checked using a Fragment Analyzer and a NGS Fragment High Sensitivity Analysis Kit. Samples were pooled equimolar, and the final pool quantified with the Kapa library quantification kit (Roche). The final pool was sequenced on a NovaSeq 6000 to produce paired-end 150 bp reads, with an average read depth of 10M reads per sample.

### RNA-seq Data Analysis

The Nextflow pipeline “rnasplice” ver 1.0.4 (https://nf-co.re/rnasplice/1.0.4/) was used to maintain ease of reproducibility in quantifying DAS. Protocol Quality control of raw reads was performed with FastQC ver. 0.12.0. Adapters were filtered with TrimGalore ver. 0.6.5. Splice-aware alignment was performed with STAR against the Arabidopsis TAIR10 ver. 59 reference genome using the default settings. Reads mapping to multiple loci in the reference genome were discarded. SAM files along with the reference genome was inputted into ASTools (49) to calculate the number of IR, ES, A3SS, and A5SS splicing events, with read length adjusted to 150 bp. Delta percent spliced in (ΔPSI) was calculated by subtracting SPHINX31 PSI by control PSI. GO enrichments for DAS events were generated by inputting up-regulated (A3SS, A5SS, IR = ΔPSI > 0.1, ES = ΔPSI < −0.05) and down-regulated (A3SS, A5SS, IR = ΔPSI < −0.1, ES = ΔPSI > 0.05) DAS events into The Ontologizer (http://ontologizer.de/; 50) with the entire transcriptome set as the background. Read depth and intron junction count visualization plots were generated using ASTool (49).

### Phosphoproteomics Sample Preparation

Total protein from 400 mg of ground tissue powder was extracted using 1:2 (w/v) extraction buffer (50 mM Tris-HCl, pH 8.0, 10 mM NA_4_P_2_O_7_, 5 mM Na_3_VO_4_, 10 mM DTT, 1% (w/v) PVPP, 2% SDS). Samples were vortexed and then boiled for 5 min at 95°C. Samples were clarified by centrifugation for 5 min at 20 000 x g at room temperature. Supernatant was transferred to new tubes and protein concentration was estimated using 1:5 sample to water for Braford quantification using Pierce™ detergent compatible Bradford assay reagent (Thermo Scientific; 1863028). Protein extracts were normalized to 1000 µg. Residues were alkylated with 30 mM iodoacetamide (IA; Sigma; I1149) and incubated for 30 min in the dark. Excess IA was quenched with 10 mM DTT for 10 min at RT. Total proteome peptides were generated using a KingFisher Apex (ThermoFisher Scientific) automated sample processing workflow as previously described without deviation (51). Peptides were digested with sequencing grade trypsin (Promega; V5113) diluted with 50 mM Triethylammonium bicarbonate (TEAB) buffer pH 8.5 (Sigma; T7408) at a ratio of 1:100 (trypsin : protein). Peptides were then acidified with trifluoroacetic acid (TFA; Fisher; A117) to a final concentration of 0.5% (v/v). Peptides were then desalted using Sep-Pak® tC18 (Waters; 186002321).

Phosphoproteome was enriched from 500 µg of total peptide using 25 µL of MagReSyn® Zr-IMAC HP bead slurry (ReSyn BioSciences; MR-ZHP005) on a KingFisher Apex. Total peptides were re-suspended in binding buffer (80% Acetonitrile; ACN, 5% TFA, and 0.1% (w/v) glycolic acid) and allowed to bind with Zr-IMAC HP beads. Bound phosphopeptides beads were washed three times in wash buffer (10% ACN + 0.2% TFA) and then eluted with 2% ammonium hydroxide (NH_4_OH). Phosphopeptides were then acidified with 100% TFA to a pH ∼2 and then desalted using OMIX C18 tips (Agilent; A5700310K) with Opentrons® OT-2 pipetting robot. C18 tips were first conditioned with one pass of 100% ACN + 0.1% TFA, followed by two passes of 60% ACN + 0.1% TFA, ending with three passes of 3% ACN + 0.1% TFA. Phosphopeptides were re-suspended in 30 µL of 3% ACN + 0.1% TFA and aspired into C18 tips. Bound phosphopetides were washed three times in 3% ACN + 0.1% TFA and then eluted in 60% ACN + 0.1%TFA.

### LC-MS/MS Analysis of Phosphopeptides

Phosphopeptides were quantified using Fusion Lumos Orbitrap mass spectrometer (Thermo Scientific) in a data dependent acquisition (DDA) mode. Phosphopeptides enriched from 500 µg of total peptide were injected using an Easy-nLC 1200 system (thermos Scientific; LC140) and separated using a 50 cm Easy-Spray PepMap C18 Column (Thermo Scientific; ES903) heated to 50°C. A spray voltage of 2.3 kV and funnel RF level of 40% was used with all data acquired in profile mode. Peptides were eluted at 300 nL/min using an upward curve gradient of solvent B [0.1% (v/v) formic acid; FA in 80% (v/v) ACN]. Separations were performed using the following 90 min solvent program: 0-18% solvent B (0-47 min), 18-28% solvent B (47-65 min), 28-46% solvent B (65-76 min), 48-100% solvent B (76-90 min). Precursor ions were measured at a resolution of 120, 000 across a mass range of 375-1550 (m/z). The full MS AGC target was set to 750% with an injection time (IT) of 20 ms. Fragment ions scans were generated using a higher-energy c-trap dissociation (HCD) fragmentation energy of 28% and measured using 15 dependent scans at a resolution of 30, 000 and a maximum IT of 60 ms. An AGC target value of 2000% and a fixed first mass of 120 m/z were used. Acquired data were searched using MaxQuant software ver 2.5.1.0 (http://www.maxquant.org/; Cox and Mann 2008) and the Arabidopsis protein database from Araport 11 (27, 533 protein encoding genes; ver. 2017-02-01; 33). Search parameters included 2 tryptic missed cleavages, Carbamidomethyl (C) fixed modification, oxidation (M) variable modification, phospho (STY) variable modification, and a peptide/protein false discovery threshold of 0.01.

### Gene Ontology Enrichment, Phosphorylation Site Visualization, and Network Figures

Significantly changing phosphosites (localization score > 0.7; q-value < 0.1) that were either up- or down-regulated (−0.58 > log_2_FC < 0.58) were inputted into The Ontologizer (http://ontologizer.de/;50) to perform a GO enrichment analysis (p-value < 0.01) with all quantified peptides as the background. GO ratios were calculated by dividing the study term by population term for each GO result. GO plots were generated using ggplot in the Tidyverse R package (52).

Phosphorylation site visualization was performed by first querying a list of splicing factors and spliceosome proteins (22) against the 2 d and 7 d phosphoproteomics data. Protein domain architecture of splicing factors was acquired from inputting protein AGI into illustrator for biological sequence (IBS) 2.0 (https://ibs.renlab.org/#/home).

Significantly changing phosphopeptides from 2 d and 7 d inhibition datasets were selected to generate association network using Cytoscape 3.10.2 (https://cytoscape.org/) with the STRING-DB ver 2.1.1. A STRING-DB score threshold of 0.65 was used to elucidate node connections. Multiple phosphosites on the same protein were averaged to generate a single log_2_FC per protein. The Cytoscape plug-in enhancedGraphics (https://apps.cytoscape.org/apps/enhancedgraphics; 53) was used to visualize log_2_FC change in phosphorylation. Protein localization information was obtained from SUBA5 (https://suba.live; 54).

## RESULTS AND DISCUSSION

### Arabidopsis Splicing-Related Protein Kinases are Conserved Across Taxa

HsSRPKs and HsCLKs have been found to regulate multiple pathways involved in cancer (32, 35, 55–57), and as such their chemical inhibition has gained traction in clinical research (33, 34, 58, 59). Highly specific and potent chemical inhibitors towards HsSRPKs and HsCLKs have passed clinical trials and are being used therapeutically. For example, Alectinib is prescribed in Canada, USA, and UK, and is known more commonly as Alecensa®. We selected four chemical inhibitors specifically designed to target splicing related protein kinases, namely HsSRPKs and HsCLKs (Supplemental Figure 2). These chemical inhibitors, such as SPHINX31, are coordinated by specific amino acid residues present within the HsSRPK active site, making them highly specific towards HsSRPKs (60) (Supplemental Figure 3). The use of splicing-related chemical inhibitors targeting the spliceosome itself have been recently adapted for plant studies, helping to answer standing questions within plant RNA splicing regulation (38, 40). However, until now, SPHINX31, Alectinib, SRPIN340, and Leucettine L41, have not been translated to plant use. Therefore, we sought to use *in silico* methods to explore the interaction between these chemical inhibitors and Arabidopsis SRPKs.

Previously, we determined that the kinase domain in each of the SRPK, AFCs/CLK, and PRP4Ks families is highly conserved across taxa (43). Correspondingly, we next aligned SRPK sequences from representative organisms of select taxa against the HsSRPKs to highlight the conserved residues and domains that perform the phosphorylation activity. We find that the SRPK bipartite domain region, boxed in blue, is highly conserved, with key amino acids maintained across taxa (Supplemental Figure 4A). Specific amino acids within the HsSRPK1 kinase domain are necessary for phosphorylating SR proteins, these include: D564, E571, D548, and K615 (61), of which all are conserved across taxa, except D548 which is E548 in *Oryza sativa (Osativ)*. Similarly, the alignment of CLKs (the Arabidopsis AFC orthologs) also found high sequence identity (Supplemental Figure 5) with the canonical LAMMER motif (L388 to R393) present in all taxa, apart from select single-celled organisms and gymnosperms (Supplemental Figure 4B). Key residues D250, R343, R346, and H382 are essential for substrate binding in the active site (62, 63), and are conserved across taxa. Finally, PRP4Ks are relatively conserved with the APE (A857 to E859) motif retained in all taxa except for *Drosophila melanogaster (Dmano)*, *Thalassiosira pseudonana* (*Tpseu)*, and *Arabidopsis thaliana*, which replaces A857 with S857 (Supplemental Figure 4C). Overall, key residues within the kinase domains of each splicing-related protein kinase are conserved across taxa, with SRPKs demonstrating a unique canonical configuration allowing for the specific inhibition by SPHINX31.

Next, we sought to determine if the amino acids targeted by the four chemical inhibitors are conserved between Human and Arabidopsis SRPKs and CLKs/AFCs. Here, we aligned HsSRPKs and AtSRPKs as well as HsCLKs and AtAFCs, focusing on the ATP binding pocket where the inhibitors are known to bind (Supplemental Figure 6). Within this multiple sequence alignment, we highlighted the residues that make close contacts with the small molecule inhibitors in the human crystal structures (PDBs: 5MY8, 4WUA, 3RAW, 5XV7). Alectinib targets eight amino acids within the HsSRPK1 phosphate binding loop (P-loop) motif of which only three of these amino acids are conserved across all AtSRPKs. This limited conservation of Alectinib binding sites in the AtSRPKs is reinforced by the higher concentrations needed to elicit a root growth effect relative to the published effective concentrations to inhibit HsSRPKs. Leucettine L41, a chemical inhibitor known to target HsCLKs (64, 65), interacts with four amino acids in the P-loop region, where all but one residue (F544) is also conserved between the AtAFCs and the HsCLKs. Similarly to Alectinib, notably higher Leucettine L41 concentrations were required to induce a root phenotypic change, suggesting this inhibitor may lack specificity towards Arabidopsis splicing-related protein kinases. Conversely, SRPIN340, a HsSRPK specific chemical inhibitor, interacts with four amino acids in the P-loop motif, which are again largely conserved in AtSRPKs. While SPHINX31, a successor inhibitor to SRPIN340, also interacts with four amino acids (L86, K109, E124, F165) within the P-loop of HsSRPK1 (60). Here, we find all targeted amino acids being largely conserved in both Group 1 and 2 AtSRPKs. In the DLG motif, all Arabidopsis SRPKs substitute A496 with V496 and L498 with F498, however, the DLG motif is canonically DFG in other protein kinases (66), and toggles between DFG-in (active) and DFG-out (inactive), essentially acting as the “activation switch” (67, 68). Although SRPIN340 and SPHINX31 target similar amino acids within the P-loop, SPHINX31 is reported to be a far more effective and specific inhibitor in comparison to SRPIN340, with SPHINX31 requiring substantially lower concentration to induce a stronger inhibitory effect (60).

### Arabidopsis Splicing-Related Protein Kinases are Structurally Conserved to Human Orthologs

With the HsSRPK specific inhibitor SPHINX31 eliciting the largest concentration dependent effect on Arabidopsis root growth, we sought to focus on Human-to-Arabidopsis SRPK structural conservation. To do this, we used Alphafold2 to model Arabidopsis SRPKs and compared them to the HsSRPK1 crystal structure (Supplemental Figure 7). We extracted predicted protein structures from Alphafold (https://alphafold.ebi.ac.uk) and highlighted regions of structural uncertainty within the model. The predicted structure of AtSRPK1 contains three regions of low confidence score (pLDDT < 70). Group I AtSRPKs have a protruding N-terminus loop (AtSRPK1: T420-G439, AtSRPK2: Q420-S440) and C-terminus loop (AtSRPK1: M1-E12, AtSRPK2: M1-E10), as well as α-helix in the middle of the protein sequence (AtSRPK1: E201-T246, AtSRPK2: T420-N244), with flexible regions on either side that sits near (∼25A) the active site and has around six positive charges all on the same face of the helix. Group II AtSRPKs have similar disordered regions, with the same C-terminus flexible region obstructing the active site (AtSRPK3: M1-D20, AtSRPK4: M1-S15, AtSRPK5: M1-S18) which protrudes further out, a disordered flexible region in the middle of the peptide sequence (AtSRPK3: T205-R292, AtSRPK4: D205-K288, AtSRPK5: D208-R303), and a longer N-terminal disordered region (AtSRPK3: P472-T538, AtSRPK4: S472-I529, AtSRPK5: G484-K534). When examining the model of HsSRPK1 we find the same regions predicted to be disordered (pLDDT < 50). The N-terminus and portion of the spacer domain disordered regions are absent from the HsSRPK1 crystal structure (69), which is to be expected since only consistently well-ordered portions of the crystal lead to observable signal. The N-terminus is thought to drive HsSRPK substrate specificity towards SR proteins, since its deletion reduces HsSRPK-to-SR protein binding affinity (70). Moreover, the disordered regions are also common across AtSRPKs and HsSRPKs families and have similar structures between Group I and Group II AtSRPKs (43). We hypothesize that these regions may also be required for substrate specificity in Arabidopsis as is found in HsSRPKs.

Next, we aligned the predicted structure of each AtSRPK to the HsSRPK1 crystal structure to determine structural conservation between both organisms (Supplemental Figure 8). Overall, the active site between each AtSRPK and HsSRPK1 is structurally similar. The N and C terminal lobes, and the spacer region which together form the active site remain in a similar position between both organisms (Figure 1C and Supplemental Figure 8). We then docked the small molecule inhibitors into the AtSRPK1 and AtSRPK4 active sites to determine the strength of interaction and predicted residues that would interact with the chemical inhibitors (Figure 1, Supplemental Figure 9-10). AtSRPK1 and AtSRPK4 were chosen as representatives since their structural alignment to HsSRPK1 had the highest RMSD value of Group I and Group II AtSRPKs, respectively. We find that the complexes with all four small molecule inhibitors have a notably low free energy (ΔG = < −9 kcal/mol) of ligand binding, indicating these inhibitors have high affinity to AtSRPKs. While the docked inhibitors occupy the active site as we expected, the orientations in the pocket are different from those observed in the crystal structures. This could be due to amino acid differences in the AtSRPKs active sites, such as F124/Y127, or the inability to account for ligand induced changes to the protein backbone and side chain position as is seen in the crystal structures of HsSRPK1 with Alectinib, SRPIN340, and SPHINX31, discussed in Batson, *et al.* (60).

**Figure 1.**
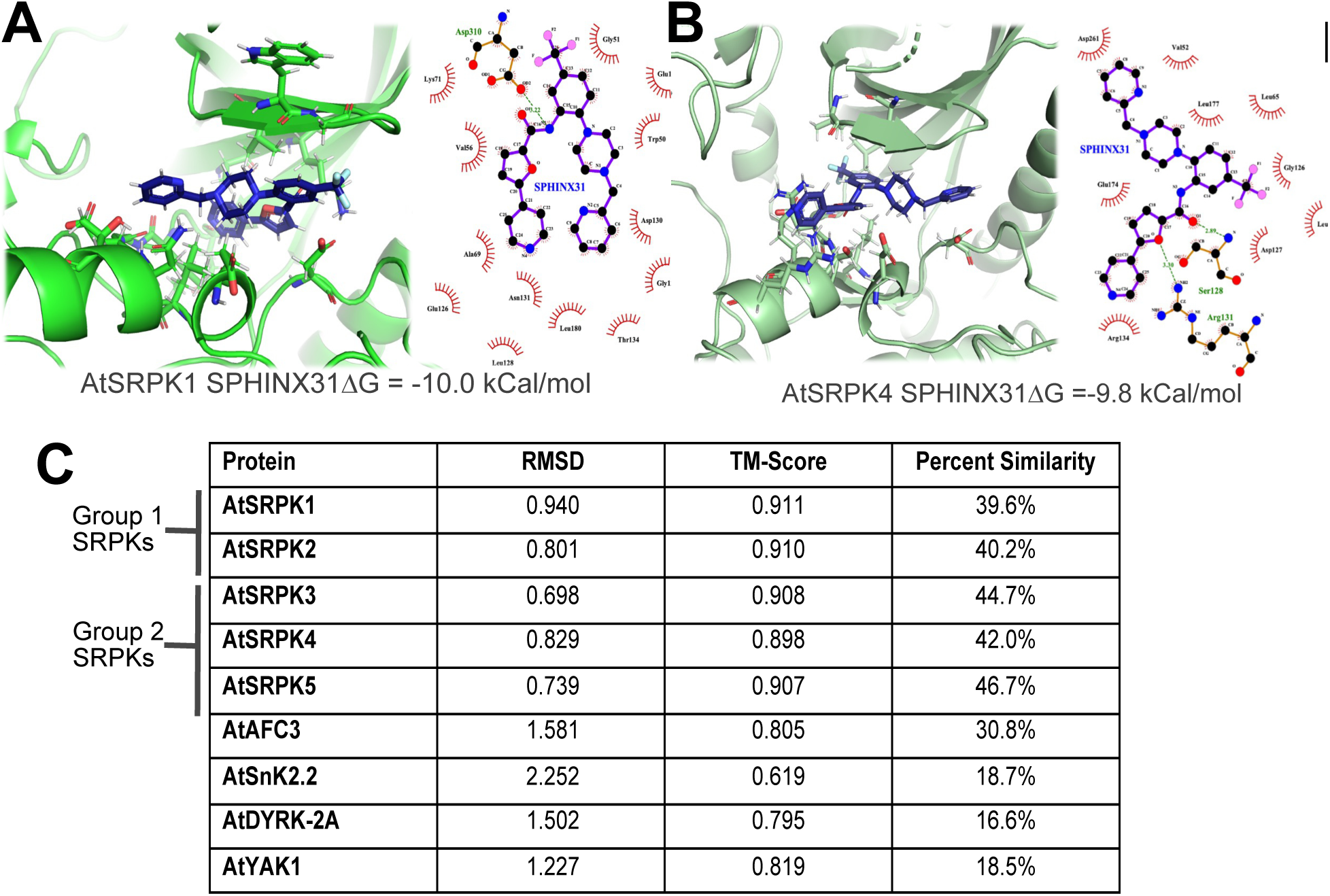
Arabidopsis AtSRPKs predicted interaction with SPHINX31 chemical inhibitor. (A) AtSRPK1 Predicted structure was imported from Alphafold and chemical structure was acquired from crystal structure PDB files (SPHINX31: 5my8). Amino acids predicted to interact with SPHINX31 were visualized using LigPlot (https://www.ebi.ac.uk/thornton-srv/software/LigPlus/). (B) AtSRPK4 interaction with SPHINX31 (C) RMSD, TM-score, and peptide sequence percent similarity between HsSRPK1 and Arabidopsis SRPK family and other protein kinases. RMSD score was calculated by aligning predicted Alphafold structures to HsSRPK1 crystal structure in Pymol. TM-Score was generated through TM-Align (https://zhanggroup.org/TM-align/). Percent similarity was generated by pairwise alignment of peptide sequences using EMBOSS needle (https://www.ebi.ac.uk/jdispatcher/psa/emboss_needle).

Finally, we sought to determine whether Arabidopsis possesses other protein kinases with structural similarities to HsSRPK1 that could potentially bind these well-known inhibitors. To do this, we queried HsSRPK1 peptide sequence against the Arabidopsis proteome using BLASTp and selected the top hits to perform peptide sequence and structural alignment. In addition to AtSRPKs and AtAFCs, we identified AtDYRK-2A (AT1G73460) and AtYAK1 (AT5G35980) as having the highest sequence similarity to HsSRPK1 (Figure 1C). However, a multiple sequence alignment (MSA) of these proteins revealed that residue identity and peptide sequence similarity are greatest between HsSRPKs and AtSRPKs (Figure 1C and Supplemental Figure 11). Next, we compared their structural similarities, since these protein kinases lack resolved crystal structures, we utilized their AlphaFold predicted structures for comparison (Supplemental Figure 12). After removing areas of low confidence (pLDDT < 70), we performed an alignment to the HsSRPK1 crystal structure. We found that both AtDYRK and AtYAK1 exhibited a lower structural similarity to HsSRPKs than AtSRPKs (Figure 1C and Supplemental Figure 13). Particularly, the spacer 1 α-helix (αS1), which is present in HsSRPKs and AtSRPKs, is absent from both AtDYRK and AtYAK1. Next, we predicted the binding of each chemical inhibitor to a completely unrelated protein kinase (SnRK2.2), to assess the potential for more wide-ranging inhibitory effects. This revealed that SPHINX31, the most specific SRPK inhibitor, does not bind the active site of either SnRK2.2, nor AtDYRK or AtYAK1 (Supplemental Figure 14). Together, these *in-silico* data suggests that SPHINX31 can specifically target Arabidopsis SRPKs.

### SPHINX31 and SRPIN340 Significantly Reduce Root Growth in Arabidopsis

Next, we grew Arabidopsis seedlings on low and high concentrations (based on applications in human studies) of either SPHINX31, SRPIN340, Leucettine L41 and Alectinib, to examine the effects of these inhibitors on Arabidopsis seedling root growth. The inhibitors with increased SRPK specificity in humans, such as SPHINX31 and SRPIN340, had significantly shorter roots compared to DMSO control throughout the entire time-course experiment (p-value < 0.0001) (Figure 2). SPHINX31, the most potent HsSRPK inhibitor (60), resulted in the most dramatic root growth reduction, with just 3 μM producing significant (p-value < 0.0001) root growth inhibition by the second day of exposure. Correspondingly, human and mice cells exposed to 3 μM SPHINX31 effectively attenuated angiogenesis and led to significant reduction in SR protein phosphorylation (71, 72). Both, SPHINX31 and SRPIN340, also had visually reduced shoot growth (not quantified). Conversely, Leucettine L41 and Alectinib, both chemical inhibitors found to target multiple human splicing-related kinases (64, 65, 73), did not have a substantial effect on root length. However, Leucettine L41 and Alectinib treated seedling showed visually longer and more abundant root hairs. However, the applied concentrations required to elicit this response is higher than typically applied in human experimentation. Given these phenotypes, we sought to quantify the number of roots hairs growing in the zone of maturation at around 2 cm above the root tip (74) (Figure 2). The differences between the SRPK specific inhibitors (SPHINX31 & SRPIN340 (60)) and the more broad-spectrum inhibitors (Alectinib & Leucettine L41 (64, 65, 73, 75)re striking when comparing root hair growth and development. SPHINX31 and SRPIN340 display a significant reduction in root hair growth. The highest concentration of SRPIN340 (10 µM) resulted in a near total abolishment of root hair formation (p-value < 0.0001) (Figure 2). A higher concentration of SPHINX31 resulted in abnormal root epidermis topography (not quantified) alongside a significant reduction in root hairs (p-value < 0.0001). The inverse occurred on seedling exposed to Alectinib and Leucettine L41, both of which resulted in a significant increase in root hair formation. While Leucettine L41 did not exhibit a dramatic increase in the number of root hairs as compared to Alecitnib, each individual root hair appeared visually longer (not quantified).

**Figure 2.**
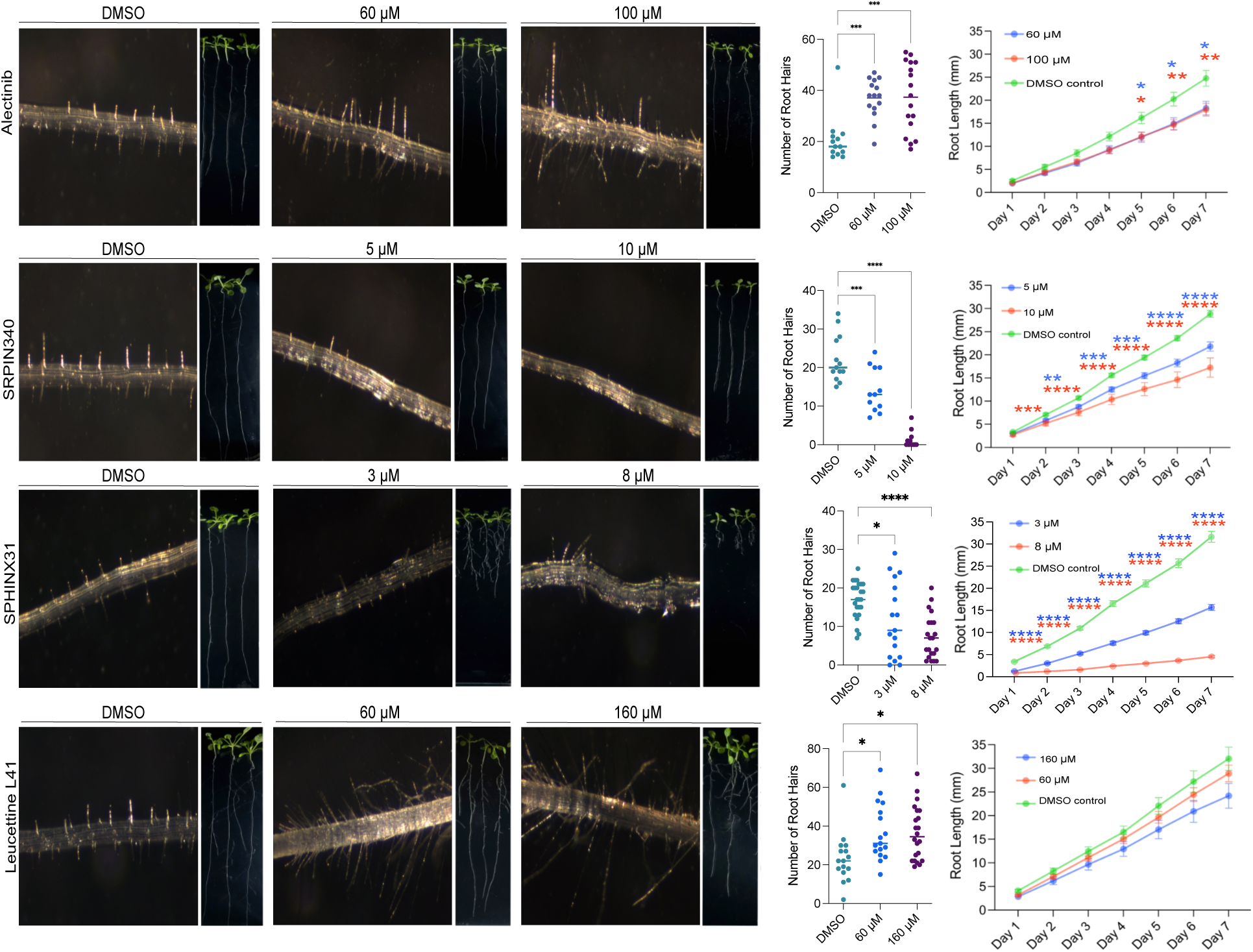
Root length and root hair number of Arabidopsis seedlings exposed to Alectinib, SRPIN340, SPHINX31, and Leucettine L41 chemical inhibitors. Seedlings were transferred from germination plates to either control (DMSO), low-, or high-concentration of inhibitor plates. Root length was measured every day for 7 d. Root hair number was quantified at 7 d of inhibition. Statistical significance was determined using a one-way ANOVA test. Asterisks represent significance level: p< 0.05= *, p< 0.01= **, p < 0.001 = ***, and p < 0.0001 = ****. Picture of seedlings and root hairs were taken at 7 d of exposure.

### Transcriptomic Analysis Reveals SPHINX31 Mediated Aberrant Splicing Across Diverse Biological Pathways

Given that SPHINX31 induced the most significant root length phenotypes at the lowest concentrations and is the most potent HsSRPK inhibitor, we next focused our molecular analyses on its short-term effects on AS by identifying and quantifying the type and number of differential alternative splicing (DAS) through RNA sequencing. To quantify aberrant splicing patterns, we calculated the differential percent spliced in (ΔPSI), which quantifies the DAS occurring in inhibited vs. control. We find a total of 3583 AS events occurring in seedlings exposed to SPHINX31 inhibition with 239 A3SS, 166 A5SS, 139 ES and 3039 IR events differentially spliced (Figure 3A and Supplemental Table 1). IR has the highest number of DAS events accounting for 85% of all AS events, while ES accounts for only 3.8% of all DAS events. Typically, IR makes up around 40-60% of all AS events occurring in the plant cell, with A5SS and A3SS forming 10-20% each, and finally ES amounting to 5-10% (76). The relatively proportions of the AS types can vary, as evidence suggests that the relative proportion of the four AS types shifts under abiotic stress, biotic stress, and across tissue types (77). Regardless of condition or tissue type, IR remains the most abundant AS type and this ratio is maintained in our data. Although A3SS and A5SS account for 11.2% of all DAS events, they exhibit the highest proportion of genes showing the most pronounced differential changes (ΔPSI of 1 or −1) (Figure 3B and Supplemental Table 1).

**Figure 3.**
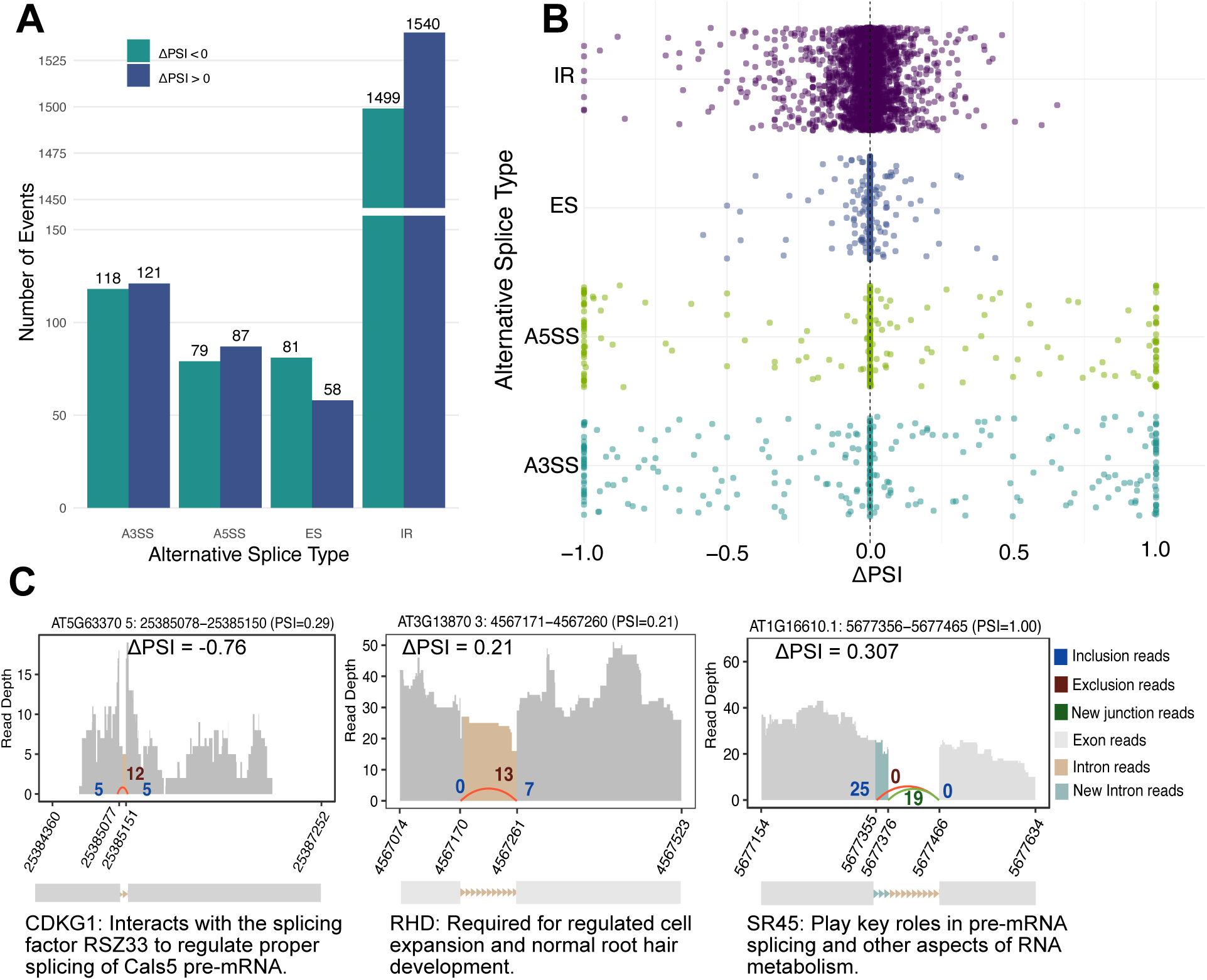
Distribution of AS events occurring in 2 d SPHINX31 inhibited seedlings. (A) number of differentially AS (DAS) events occurring under each splice type (ΔPSI > 0 = occurring more in SPHINX31, ΔPSI < 0 = occurring less in SPHINX31 compared to WT). (B) Distribution of DAS intron retention (IR), exons skipping (ES), alternative 5’SS (A5SS), and alternative 3’ SS (A3SS) occurring in SPHINX31 inhibited tissue. (C) Visualization of the number of reads mapped to the intron retention event detected by ASTool. TAIR gene name and descriptions are positioned below depiction of the intron retention. Differential PSI for each gene is denoted by ΔPSI which the PSI of SPHINX31 treatment minus PSI of control. Inclusion reads are the mapped reads that include intron in the mRNA, while exclusion reads are the mapped read that exclude the intron in the mRNA. New junction reads are the number of mapped reads that denote portion of intron included in the mRNA. Grey bars represent the number of reads mapped to the exon and beige bars represent the number of reads mapped to the intron. Teal bars represent number of read mapping to a portion of the intron, termed new intron.

We then performed GO enrichment analysis for biological processes to get insight into which biological pathways are associated with DAS genes in SPHINX31 inhibited tissue. We find genes with up-regulated A3SS splicing patterns (ΔPSI > 0.1) are associated with DNA remodelling, light sensing, among others (Supplemental Figure 15; Supplemental Table 2). While a down-regulation of A3SS events are related to N-terminal protein amino acid modification, mRNA processing, protein assembly, among others. Similarly, SPHINX31 causes a down-regulation of A5SS splicing pattern in genes with biological GO terms related to mRNA processing, lateral root formation, regulation of mRNA export from nucleus, among others (Supplemental Figure 16 and Supplemental Table 2). In Supplemental Figure 17, we find that SPHINX31 causes higher differential ES of genes related to stem cell differentiation, RNA processing, chloroplast-nucleus signalling pathway, among others (Supplemental Figure 16 and Supplemental Table 2). Finally, down-regulated IR events had a significant enrichment in GO terms related to mRNA transport including export from the nucleus and nucleocytoplasmic transport (Supplemental Figure 18 and Supplemental Table 2). There was also an enrichment of terms related to root development and cell differentiation such as root system development, regulation of post−embryonic root development, and stem cell division. Together, this suggests that dysregulated AS, particularly aberrant IR, likely contributes to the root phenotypes exhibited under SPHINX31 inhibition.

### SPHINX31 Induces Aberrant Splicing of RNA Splicing-Related Proteins

We find numerous splicing factors with DAS patterns in seedlings treated with SPHINX31 inhibitor. Of particular note was the substantial decrease in A3SS of *ARABIDOPSIS SR-LIKE 1* (*ATSRL1*; AT5G37370; ΔPSI = −1), *SERINE ARGININE RICH 45* (*SR45;* AT1G16610; ΔPSI = −0.6), *ARABIDOPSIS FUS3-COMPLEMENT GENE 2* (*AFC2*; AT4G24740; ΔPSI = −0.3) (Supplemental Table 1). Splicing factors with down-regulation of A5SS include *SR45* (ΔPSI = −1), *SPLICING FACTOR 3B* (*SF3B5B*; AT4G14342; ΔPSI = −1), *CC1-LIKE SPLICING FACOTR* (*CC1*; AT2G16940; ΔPSI = −1), *RNA-BINDIGN PROTEIN 47A* (*RBP47A*, AT1G49600, ΔPSI = −0.94), and *ARGININE/SERINE-RICH ZINC KNUCKLE-CONTAINING PROTEIN 33* (*RS2Z33*; AT2G37340; ΔPSI = −1). There are also some splicing related genes with down-regulated IR such as *CYCLIN-DEPENDENT KINASE G1* (*CDKG1*; AT5G63370, ΔPSI = −0.71), and to a lesser degree *ARABIDOPSIS THALIANA ORTHOLOG OF HUMAN SPLICING FACTOR SC35* (*SC35;* AT5G64200; ΔPSI = −0.115), and *SERINE ARGININE SPLICING FACOTR 34* (*SR34*; AT1G02840; ΔPSI = −0.104). Arabidopsis SR transcripts are known to undergo extensive AS themselves, with their splicing patterns varying in response to environmental stress (78, 79) and during development (80). The AS of splicing factors not only regulates their own functions but also influences the splicing patterns of other SR splicing factors (81, 82), forming an immensely complex regulatory network. In our data, we see various splicing factors that are implicated in the regulation of AS of other splicing factors. For example, *SR45* has seven splice variant isoforms, of which two have been characterized as playing distinct roles in plant development (80). The constitutively spliced variant, *SR45.1*, contains a potential phosphorylation site at S219 and is essential for normal root growth. While *SR45.2* is produced through A3SS that removes 21-nucleotide sequence containing S219 site (80, 83). Thus, it is possible that SPHINX31 interferes with AtSRPK ability to regulate AS pathways that drive the production of *SR45.2,* leading to irregular root growth. Additionally, the inhibition of AtSRPKs may prevent the AS of splicing factors mediated by SR45. For example, a loss of *SR45* results in aberrant splicing of other SR proteins (84), including altered A5SS of *CC1* (*85*); an AS event seen in our data.

SPHINX31 also causes high exclusion of A5SS pattern in *SF3B5B* (ΔPSI = −1) (Supplemental Table 1). Human SF3B5B associates with other splicing proteins to form the U2 snRNP complex (86, 87). Very little is known about SF3B5B in plants, however Ishizawa et al., (2019) described a novel Arabidopsis mutant line with enhanced root hair formation in response to light (*light-sensitive root hair development 1; lrh1)*(*88*). This work revealed protein p14, a putative component of SF3B, as being responsible for the *lrh1* phenotype. Application of the spliceosome inhibitor PB mimicked the *lhr1* root phenotype, suggesting AS is a key mechanism in regulating root hair development in response to light signals. It is thus possible that inhibiting AtSRPKs with SPHINX31 causes dysregulating of splicing signals that then result in aberrant splicing of genes involved in root hair formation and light-sensing, both of which are enriched GO terms in our DAS analysis (Supplemental Table 2).

SPHINX31 also induces aberrant IR in *CDKG1* (ΔPSI = −0.71), which is a component of the recombination and pairing machinery needed for meiotic and somatic division (89). *CDKG1* is alternatively spliced with a longer IR isoform containing a nuclear localized signal and is named *CDKG1L*, while *CDKG1S* is the short isoform without the intron. CDKG1S lacks two out of four SR domains and the NLS sequestering the protein in the cytoplasm (90) CDKG1 contains a RS domain that interacts with RSZ33 to regulate the proper splicing of *CALLOSE SYNTHASE* (*CALS5),* and thus pollen wall formation (91). Huang et al., (2013) propose RSZ33 recruits CDKG1 to the U1 snRNP to facilitate the splicing of *CALS5* which induces proper pollen development, more direct evidence is required to validate this pathway. Under normal conditions *CDKG1* IR has a PSI of 0.8 (PlaASDB; http://zzdlab.com/PlaASDB/ASDB/search.php), while SPHINX31 exposure results in the CDKG1 intron to be excised at a higher level (ΔPSI = −0.71), suggesting inhibition of splicing-related protein kinases dysregulate the splicing of *CDKG1* and consequently affect downstream splicing of genes that *CDKG1* would normally splice. Thus, it is possible that inhibiting AtSRPKs triggers a cascade of abnormal AS events, where splicing factors themselves are incorrectly spliced, affecting their ability to mediate spliceosome splice site selection, leading to further mis-splicing of other splicing factors.

### SPHINX31 Inhibition Dysregulates the Alternative Splicing of Cell Division, Auxin-, and Root Development-Related Transcripts

#### Root Meristem

There are two splice variants of *P23-1* (AT4G02450) one of which includes an A3SS event on the primary exon that removes a 225 bp region. Under SPHINX31 inhibition, we find a dramatic increase in A3SS patterns of *P23* (ΔPSI = 1) (Supplemental Table 2). P23-1, alongside P23-2, form co-chaperones of HEAT SHOCK PROTEIN 90 (HSP90) (92), and loss of *P23-1* results in short roots and reduced meristem lengths (93). Additionally, *P23-1* mutants exhibit decreased expression of PIN1 and PIN7, leading to inappropriate distribution of auxin and impaired root elongation. The function of the A3SS splice variant of *P23-1* remains uncharacterized in plants, however, its canonical spliceform is known to be phosphorylated and has a role in certain signalling pathways. Canonical P23-1 is phosphorylated by Ser/Thr protein kinase CASEIN KINASE 2 (CK2) and this phosphorylation event is necessary for normal root development, with CK2 phosphoablative expressing lines showing lower root meristem cell number and shorter roots (94).

Further, meristem development requires cell-to-cell trafficking, rendering it possible that SPHINX31 may be preventing AS patterns required for proper production of trafficking related proteins. For example, we find multiple genes in the ARF GTPase family genes displayed dramatic DAS A5SS patterns including *ADP-RIBOSYLATION FACTOR A1F* (*ARFA1F*; AT1G10630; ΔPSI = −1), *ADP*-*RIBOSYLATION FACTOR-LIKE A1C* (*ARLA1C*; AT3G49870; ΔPSI = −1), *ADP*-*RIBOSYLATION FACTOR-LIKE A1D* (*ARFA1D*; AT1G70490; ΔPSI = −0.994). The ARF family of proteins are involved in membrane trafficking steps and the regulation of vesicle trafficking (95, 96). In Arabidopsis ARFA1s function in vesicle trafficking and their function is essential for proper pollen development (97). Although the different splice variants of *ARF* have not been studied it is possible that their dysregulated AS patterns as a result of SPHINX31 treatment is preventing proper root and meristem development through impaired vesicle trafficking.

#### Auxin

We also observed an increase in the A5SS splicing pattern of *PATATIN-RELATED PHOSPHOLIPASE A* (*PLP1*; AT4G37070) also known as *PHOSPHOLIPASE A IVA* (*PLAIVA*) (ΔPSI = 0.99)(Supplemental Table 2). This enzyme cleaves phospho- and galactolipids to generate free fatty acids and lysolipids that are required for plant hormone signalling (98). There are four *PLP1* isoforms generated through IR, A5S, or both, though their functions have not been reported. PLAIVA is expressed strongly in the root and removal of the protein results in reduced lateral root development, reminiscent of impaired auxin response (99). *In vitro*, CPK3 phosphorylates the PLAIVA protein, enhancing substrate selectivity towards phosphatidylcholine, the main source of auxin-induced fatty acid release. Both, canonical P23-1 and PLAIVA are regulated at the PTM level, however, they have a higher proportion of DAS transcripts under SPHINX31 treatment, suggesting that these auxin-related proteins may also be post-transcriptionally regulated and their AS patterns inducing root phenotypes.

Moreover, we find an *AUXIN RESPONSE FACTOR 2* (*ARF2*; AT5G62000) with down-regulated A3SS splicing pattern under SPHINX31 inhibition (ΔPSI = −0.3). *ARF2* has five splice variants generated from a variety of AS events; IR, A5SS and A3SS. *ARF2* is a member of the ARF family of transcription factors known to regulate auxin (100). Not much is known about the AS variants of *ARF2,* however, a study in Tobacco found that *ARF2* is differentially spliced in the root under K^+^ stress (101), suggesting that there are biological functions to the AS of *ARF2*. ARF2 is a negative regulator acting downstream of xyloglucan-dependent control of hook development and transcription control of polar auxin (102). Plants lacking *ARF2* have irregular distribution of PIN proteins and therefore auxin, which affects primary root growth (103). Moreover, ARF2 negatively regulates PLETHORA 2 (PLT2) at the protein level in root meristems, and lower activity of PLT2 is associated with reduced cell division and inhibits cell differentiation in root meristems. The combination of high differential AS of auxin-related genes and GO enrichment terms for auxin suggests that the root phenotypes under SPHINX31 could be due to altered AS patterns of genes that regulate auxin and thus inhibit proper root development. Moreover, many of the most dramatic DAS belong to genes that are involved in auxin distribution, root elongation, RNA splicing, and meristem function, suggesting that SPHINX31 may be preventing AtSRPK’s ability at regulating AS pathways critical for auxin signalling, cell trafficking, and stress regulation, ultimately leading to impaired root development.

#### Root development

*ECTOPIC ROOT HAIR 1* (*ERH1*; AT1G05850), also known as *CHINATAE-LIKE PROTEIN1* (*CTL1*) encodes for two splice variants of which one is formed through A5SS. CTL1 is essential for salt, heat, and drought stress, and is involved in root hair development (104–106). Mutations to *CTL1* causes altered cell wall composition and decreased abiotic stress resistance due to reduced cellulose content (105, 107). CTL1 is highly expressed in the roots, particularly in the cell wall of roots, and mutation to *CTL1* resulted in less crystalline cellulose and alteration to pectin formation which led to abnormal root development (108). We find A5SS AS of *CTL1* is dramatically decreased under SPHINX31 inhibition (ΔPSI = −1). Moreover, SPHINX31 caused irregular formation of the root epidermis (Figure 2), suggesting that genes, like *CTL1* are irregularly spliced and their AS patterns affect root epidermis development.

### SPHINX31 Treatment Drives Down-Regulation of Phosphosites Linked to Reproduction, Cytoskeleton, and RNA Splicing

Arabidopsis SRPKs phosphorylate SR dipeptide repeats on splicing factors, which can alter their activity (109, 110), subcellular localization (111), and can subsequently affect AS patterns of downstream genes (112). Thus, we next sought to determine changes in the phosphorylation landscape as of result of AtSRPK inhibition. Here, we conducted a phosphoproteomic analysis following 2 and 7 d of SPHINX31 exposure, quantifying a total of 5792 phosphosites (Figure 4A and Supplemental Table 3). Of these, 799 significantly changing phosphosites were quantified after day 2 and 409 phosphosites after day 7 of exposure (FDR < 0.1, localization score > 0.7). At 2 d of inhibition, 148 phosphosites were significantly down-regulated (log_2_FC < −0.58), with some of the most dramatically changing phosphosites linked to RNA splicing, such as SCARECROW-LIKE TRANSCRIPTION FACTOR 11 (SCL11; AT5G59460), ARGININE/SERINE-RICH SPLICING FACTOR 31 (RS31; AT3G61860), OVATE FAMILY PROTEIN 5 (OFP5; AT4G18830) among others (Figure 4B, 4D, and Supplemental Table 4). We also find hnRNPs and other spliceosome proteins with significant down-regulated phosphosites, such as COLD CIRCADIAN RHYTHM AND RNA BINDING 2 hnRNP (CCR2; AT2G21660; 2 d log_2_FC = −3.62), an ortholog to human hnRNP A/B (113) (Supplemental Table 3 and 5). Other significantly changing phosphosites were related to embryogenesis, such as BUD SITE SELECTION PROTEIN 13 (BUD13; AT1G31870) whose function is necessary for the splicing of genes involved in early embryogenesis and root meristem development (114, 115). While others, like OFP5, are involved in cytoskeleton organization (116). Phosphosites that were down-regulated at 2 d had an enrichment of biological GO terms related to reproduction, protein complex disassembly, cell cycle and cytoskeleton, amongst other (Figure 4C and Supplemental Table 5).

**Figure 4.**
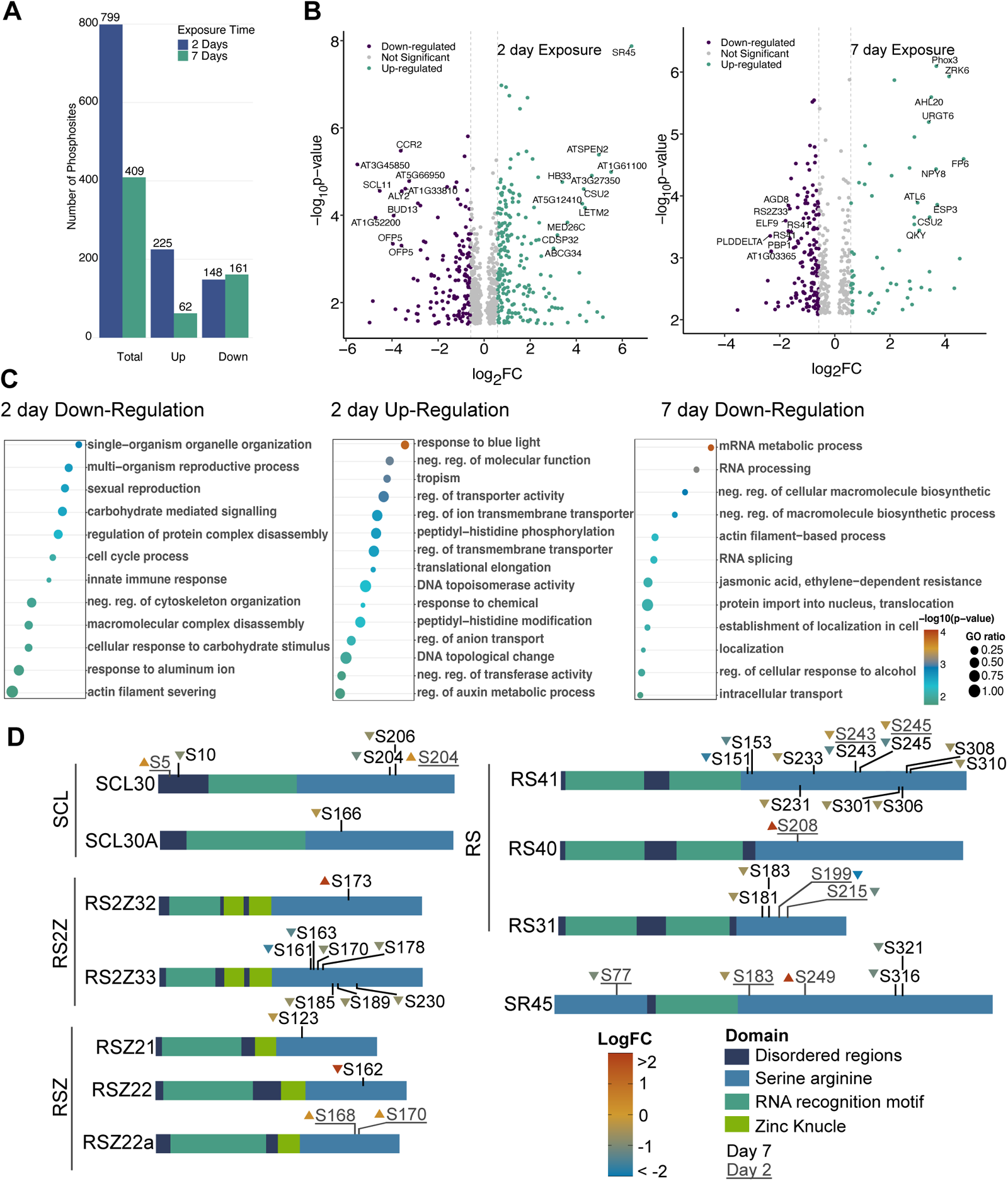
Analysis of quantitative phosphoproteomic of 2 d and 7 d SPHINX31 inhibited Arabidopsis seedlings. (A) total number of up- and down-regulated phosphosites. (B) Volcano plot of significant changing phosphosites identified. Most significantly changing phosphosites were labelled with their TAIR protein name. (C) Gene ontology enrichment of significantly changing phosphosites. significant (p-value < 0.01, down-regulated: logFC < −0.58, up-regulated: logFC > 0.58. No significant GO terms were found for 7d up- regulated phosphosites. (D) Significantly changing phosphosites identified on SR splicing factors after 2 d (underscored) and 7 d of SPHINX31 inhibition.

Conversely, we find other phosphosites on RNA splicing related proteins that are up-regulated (log_2_FC > 0.58), such as SERINE ARGININE SPLICING FACTOR 45 (SR45; AT1G16610), SERINE ARGININE SPLICING FACTOR 40 (SR40; AT4G25500), COP1 SUPPRESOR 2 (CSU2; AT1G02330) (Figure 4B). Suggesting that by 2 d, the plant may have compensated for the loss of AtSRPK activity by activating alternative phosphorylation pathways targeting these splicing factors. Alternatively, it is also possible that AtSRPK inhibition prevents the activation of proteins responsible for the dephosphorylated status of SR45, for example. After 7 d of growth in SPHINX31 plates, some of the most strongly down-regulated phosphosites also belong to RNA splicing factors such as SERINE ARGININE RICH 41 (SR41; AT5G52040; log_2_FC = −1.68), RS2ZZ33 (AT2G37340; log_2_FC = −1.6), SERINE ARGININE 22 (SR22; AT4G31580; log_2_FC = −2.50), SC35-LIKE 30 (SCL30; AT3G55460; log_2_FC = −1.13)(Figure 4D, Supplemental Table 3, and Supplemental Table 4). Down-regulated GO terms were found to be associated with RNA processing, RNA splicing, cytoskeleton, localisation, protein import into the nucleus and other transport related terms (Figure 4C and Supplemental Table 5). The continued reduction in root growth up to day 7 of SPHINX31 exposure, coupled with the down-regulation of RNA splicing protein phosphorylation, indicates that AtSRPK inhibition cannot be fully circumvented, highlighting their essential role in root development. Further analyses are needed to determine whether the AtSRPK-mediated root development phenotype arises from dysregulated AS of genes involved in the auxin signalling pathway.

### SPHINX31 Induces Up-Regulation of Phosphosites Linked to Light Signalling and Photomorphogenesis

Up-regulated phosphosites following 2 d of SPHINX31 treatment are significantly enriched in GO terms related to blue light response, tropism, and auxin metabolism, all biological processes central to photomorphogenesis. AS, protein degradation, and photomorphogenesis are intricately linked (117). For example, spliceosome degradation and IR of photomorphogenic genes are critical for light-induced seedling morphogenesis (3). The authors found that, in response to light, the E3 ubiquitin ligase CONSTITUTIVE PHOTOMORPHOGENIC 1 (COP1) ubiquitinates the spliceosome component DOMNIANT COP1-6 SUPPRESOR 1 (DCS1), which alters the confirmation and activity of the spliceosome. This modification promotes the IR of negative regulators of light signalling pathway, sequestering these transcripts in the nucleus and preventing their translation, thereby facilitating seedling photomorphogenesis. COP1 activity must be tightly controlled to ensure proper protein accumulation of its target proteins in response to environmental cues. CSU2, a key regulator, directly interacts with COP1 in nuclear speckles, supressing its activity and preventing degradation of photomorphogenesis-related proteins, particularly those affecting primary root growth (118). In our data, we find substantial increases in CSU2 phosphorylation (2 d log_2_FC = 4.32 and 7 d log_2_FC = 3.44). While CSU2 phosphorylation is annotated in publicly available databases (e.g. PTMviewer), its functional significance remains unexplored. However, it is possible phosphorylated CSU2 is affecting COP1 activity. Additionally, COP1 regulates shoot-to-root auxin transport by controlling PIN1 and PIN2 proteins (119), whereby auxin produced in the shoots under illuminated conditions is transported to the roots via PIN proteins, promoting primary root development (120, 121). Conversely, under shade or darkness, root growth is stunted. Our data suggests that SPHINX31 inhibition of SRPKs may disrupt light signalling pathways, impairing auxin transport from the shoots to the roots. Disrupted auxin transport can lead to shorter roots, mimicking the phenotypes of dark grown plants. Light sensing disruption is supported by the up-regulation of phosphosites on photoreceptors of key photoreceptors and associated proteins, including PHOTOTROPIN 1 (PHOT1; AT3G45780; log_2_FC = 0.74), PHYTOCHROME B (PHYB; AT2G18790; log_2_FC = 0.83), and the phytochrome-interacting proteins FKBP12 INTERACTING PROTEIN 37 (FIP37; AT3G54170; log_2_FC = 1.49). Furthermore, we find direct repressors of COP1, such as CSU2 and PIF4 (log_2_FC = 1.01), to have highly up-regulated phosphorylation statuses, potentially functioning to alter COP1 activity, and subsequently affecting PIN proteins, auxin transport, and/or the AS of other photomorphogenic genes. Further studies are needed to uncover how CSU2 phosphorylation may impact COP1 activity, auxin transport, and root development in the context of photomorphogenesis, as well as to position AtSRPKs within these pathways.

### Inhibiting AtSRPKs Unveils a Complex Interplay Between Phosphorylation and Alternative Splicing

Overall, our analyses reveal numerous splicing factors that exhibit both DAS and changes in their phosphorylation status in response to SPHINX31 exposure (Figure 4B, Figure 5, Supplemental Table 1, and Supplemental Table 3). Splicing factors are subject to multiple layers of regulation, including PTMs and AS. For instance, SR45 is not only regulated by PTMs but also undergoes extensive AS, both of which play crucial roles in its functional dynamics. At the PTM-level, SR45 phosphorylation on T264 influences its ABA-induced accumulation (112). SR45 phosphorylation also induces its translocation to nuclear speckles allowing it to interact with AFC3 *in vitro* (122). Recently, it has been shown that Group II AtSRPKs phosphorylate SR45, promoting its association with the ASAP complex subsequently leading to the expression and AS of flowering genes (37). At the post-transcriptional level, *SR45* is alternatively spliced to produce *SR45.1* the isoform that is required for a root development (123). The AS of *SR45* is mediated by other splicing factors, some of which, are activated by phosphorylation events. Our data indicate that several splicing factors experience down-regulation of their phosphorylation status along with dramatic DAS patterns under SPHINX31 treatment, such as is observed with SR45 (figure 4B and 4D). It remains challenging to determine whether the detected DAS patterns are a direct consequence of AtSRPK inactivation or the downstream effects of disrupted AtSRPK-mediated phosphorylation, as the interplay between PTMs and AS relating to splicing factors like SR45 highlights the complexity of post-transcriptional regulation and the broader implications of AS dysregulation.

**Figure 5.**
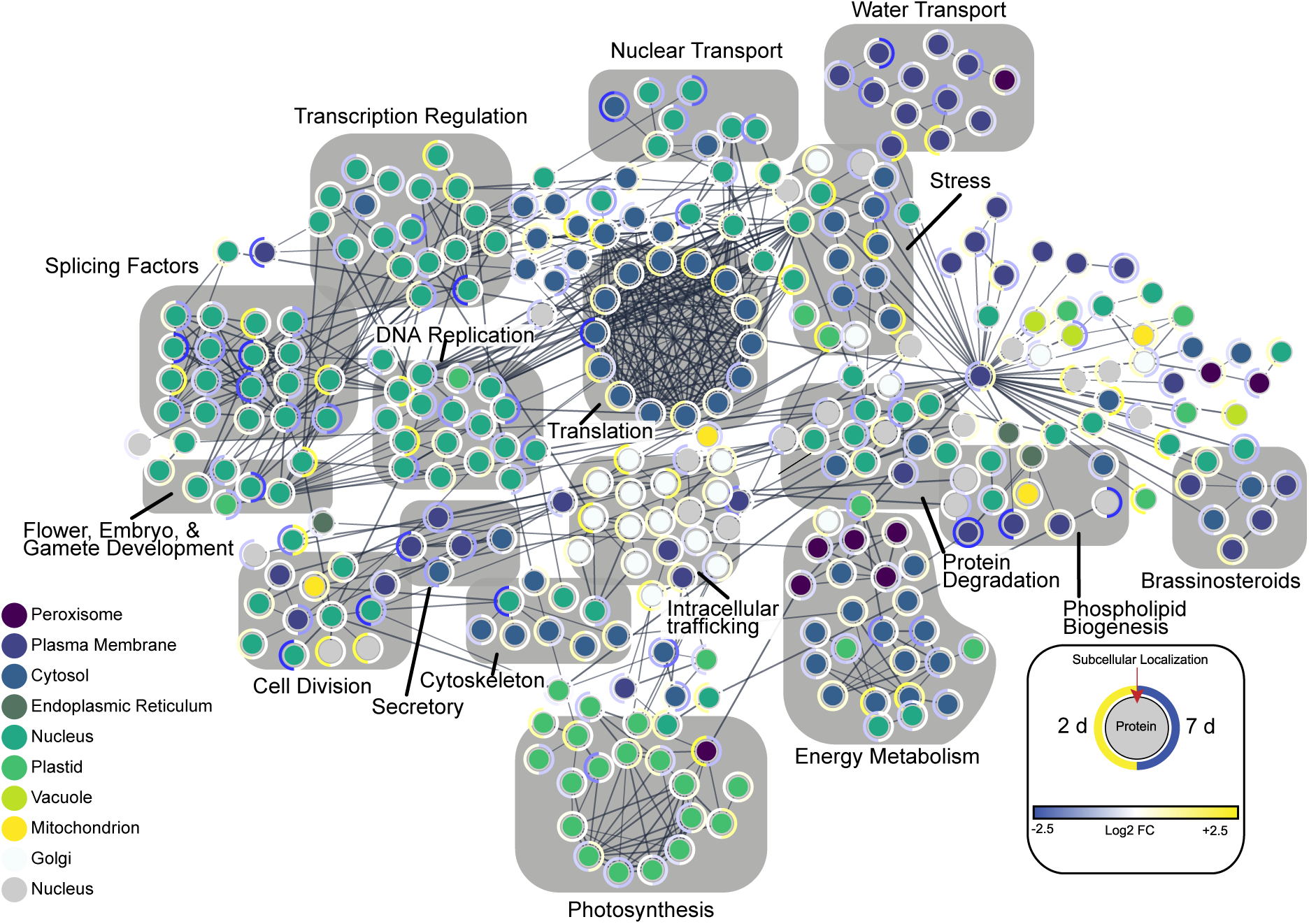
Association network of significantly changing phosphosites after 2 d and 7 d SPHINX31 inhibition. Edge thickness represents STRING-DB calculated edge score of 0.65. Inner circle represents sub-cellular location annotated by SUBA5. Outer ring of nodes represents logFC values, with left side representing 2 d and right side representing 7 d value. Grey blocks represent groups of nodes belonging to similar biological processes.

## Conclusion

The pivotal role of splicing-related protein kinases, such as SRPKs, is well appreciated in clinical research, with numerous chemical inhibitors developed over the years to block their enzymatic activity. Many of these inhibitors are used in cancer therapies, where they prevent HsSRPK from activating splicing factors that drive AS of cancer inducing genes (32, 56, 59). Despite the extensive use of protein inhibitor molecules in plant research, including protein kinase inhibitors (e.g. rapamycin), only recently have human-designed spliceosome inhibitors been applied in plant studies as a method of investigating plant RNA splicing (3, 38). To date, none have used human-designed chemical inhibitors that target the canonical plant protein kinases that transmit signals to the spliceosome. In this study, we sought to use four chemical inhibitors, SPHINX31, SRPIN340, Leucettine L41, and Alectinib to explore the effects of inhibiting Arabidopsis splicing-related protein kinases. We find that the most specific SRPK chemical inhibitors, SPHINX31 and SRPIN340, caused the most significant reduction in root growth and almost completely abolished root hair formation. We then focused our efforts on SPHINX31, the most potent inhibitor tested, quantifying differentially alternative spliced genes and phosphosites resulting from its application. Our data suggests that AtSRPKs regulate multiple AS events involved in root development, particularly biological processes related to meristem differentiation, cytoskeletal organization, and auxin distribution; all processes that are interconnected and involved in root growth and development (Figure 6). Overall, our use of SPHINX31 has revealed substantial new knowledge of the roles AS plays in plants, in addition to new fundamental insights about the PTM-level regulation of AS.

**Figure 6.**
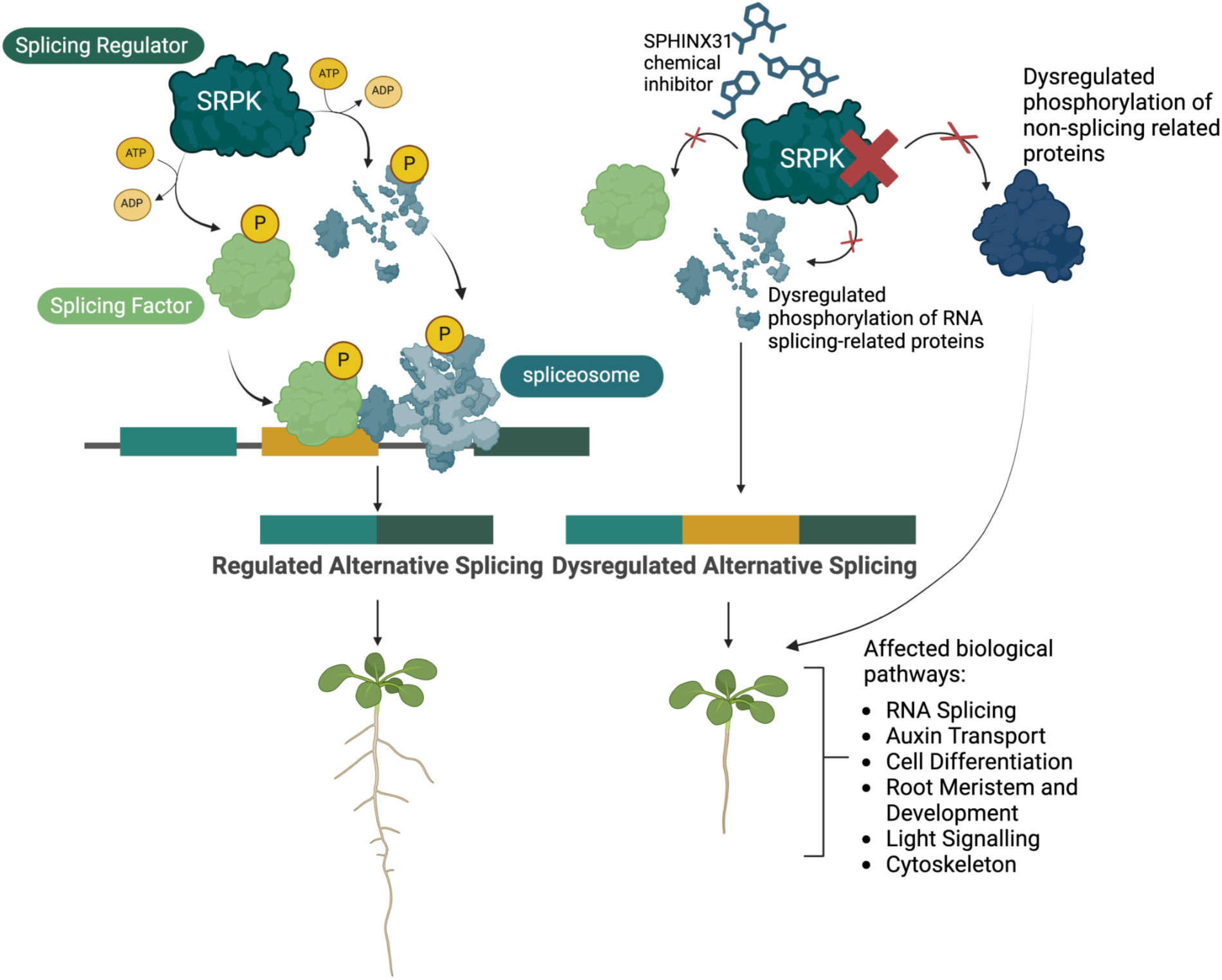
Schematic of the proposed mechanism illustrating the effects of inhibiting Arabidopsis SRPKs with SPHINX31.

## ACKNOWLEDGEMENTS

The authors would like to thank Dr. Jack Moore from the Alberta Proteomics and Mass-Spectrometry Facility for technical support and Dr. Rhiannon Peery for helpful discussion on RNA sequencing data analysis. We would also like to thank Annelien Verfaillie, Khaled Mirzai, and Alvaro Cortes Calabuig from the GenomicsCore KU Leuven for technical support.

## AUTHOR CONTRIBUTIONS

*Maria Camila Rodriguez Gallo*: Investigation, Conceptualization, Data curation, Formal analysis, Visualization, Methodology, Writing—original draft. *Cameron Murray*: Writing—review & editing, Formal analysis, Visualization. *Curtis Kennedy*: Formal analysis, Data curation. *Mohana Talasila*: Investigation. *Devansh Bhatt*: Investigation. *Joselyn Steyer*: Investigation. *Mark Glover:* Supervision. Writing— review & editing. *R. Glen Uhrig*: Supervision, Conceptualization, Writing—review & editing, Funding acquisition.

## SUPPLEMENTARY DATA

Supplementary Data are available online.

## DATA AVAILABILITY

All raw proteomics data can be found through the Proteomics IDEntifications Database (PRIDE) using the dataset identifier PXD060930. RNA sequencing data has been deposited in the European Nucleotide Archive (ENA) at EMBL-EBI under accession number PRJEB85743.

## SUPPLEMENTAL FIGURES

**Supplemental Figure 1. Depiction of imaged area for root hair quantification.**

**Supplemental Figure 2. Structure of the four chemical inhibitors applied to Arabidopsis seedlings.**

**Supplemental Figure 3. Crystal structure of HsSRPK1 with SPHINX31 in the active site.**

**Supplemental Figure 4. Peptide alignment and percent conservation of human splicing-related protein kinases to orthologs from various species.**

**Supplemental Figure 5. AtAFCs protein sequence alignment percent identity comparisons to HsCLKs.**

**Supplemental Figure 6. Protein sequence alignment of HsSRPK to AtSRPK to show amino acid conservation targeted by each of the four chemical inhibitors.**

**Supplemental Figure 7. AtSRPK and HsSRPK 3-dimensional structure visualized with Alphafold confidence scores.**

**Supplemental Figure 8. AtSRPK family superimposed on HsSRPK1 highlighting structural confirmation.**

**Supplemental Figure 9. AtSRPK1 predicted interaction with the four chemical inhibitors.**

**Supplemental Figure 10. AtSRPK4 predicted interaction with the four chemical inhibitors.**

**Supplemental Figure 11. Protein sequence alignment of HsSRPKs and Arabidopsis splicing-related kinases (AtSRPKs and AtAFCs), alongside other Arabidopsis proteins.**

**Supplemental Figure 12. Other Arabidopsis proteins 3-dimensional structure visualized with Alphafold confidence scores.**

**Supplemental Figure 13. Other Arabidopsis proteins aligned on HsSRPK1 highlighting structural confirmation.**

**Supplemental Figure 14. AtSnRK2.2 predicted interaction with SPHINX31.**

**Supplemental Figure 15. Biological gene ontology enrichment of A3SS alternative spliced genes in SPHINX31 inhibited Arabidopsis seedlings.**

**Supplemental Figure 16. Gene ontology enrichment of A5SS alternative spliced genes in SPHINX31.**

**Supplemental Figure 17. Gene ontology enrichment of ES alternative spliced genes in SPHINX31.**

**Supplemental Figure 18. Gene ontology enrichment of IR alternative spliced genes in SPHINX31.**

## SUPPLEMENTAL TABLES

**Supplemental Table 1. Alternative splicing quantification using ASTools.**

**Supplemental Table 2. GO enrichment results of alternative splicing results.**

**Supplemental Table 3. 2 d and 7 d SPHINX31 inhibition phosphoproteomics data.**

**Supplemental Table 4. GO enrichment results of significantly changing 2 d and 7 d phosphosites.**

**Supplemental Table 5. Phosphosites of splicing related proteins.**

