## Supplemental Figures for "Chemical Inhibition of Splicing-Related Protein Kinases Reveals Phosphorylation-Driven Regulation of RNA Alternative Splicing in *Arabidopsis* Seedlings"

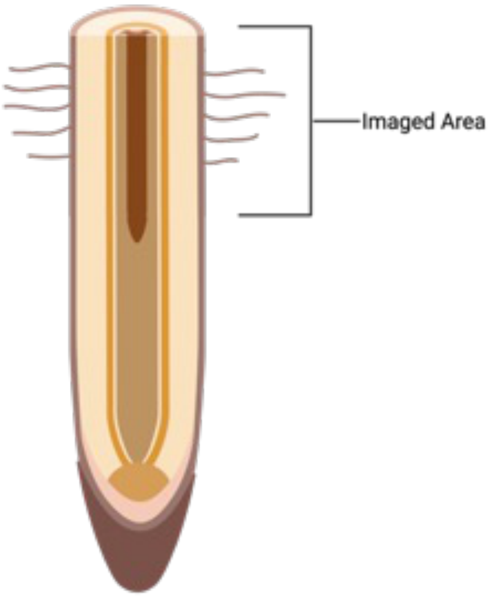


***Supplemental Figure 1. Depiction of imaged area for root hair quantification.*** *Pictures were taken approximately 2 cm above the root tip corresponding to the zone of maturation.*

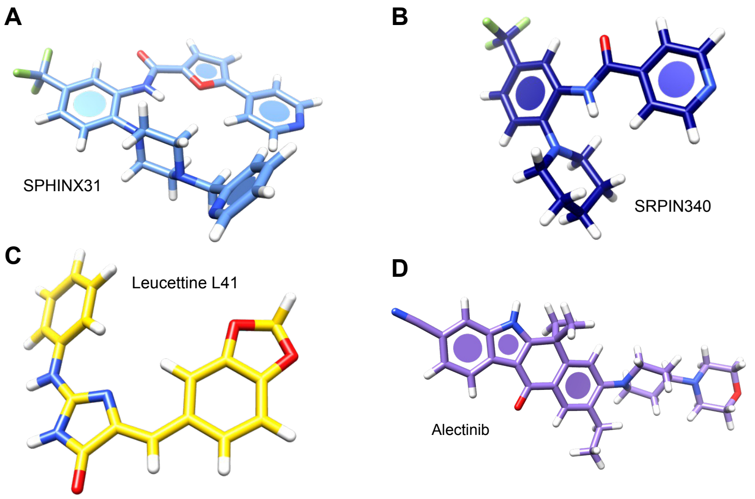


***Supplemental Figure 2. Structure of the four chemical inhibitors applied to Arabidopsis seedlings.*** *Images were generated in PyMol vers 3.1 (https://www.pymol.org) by importing the crystal structure from protein data bank (PDB) (https://www.rcsb.org) into Pymol vers 3.1. (a) SPHINX31, PDB code: 5my8 (b) SPRIN340, PDB code: 4wua (c) Leucettine L41, PDB code: 8P05 (d) Alectinib, PDB code:5XV7.*

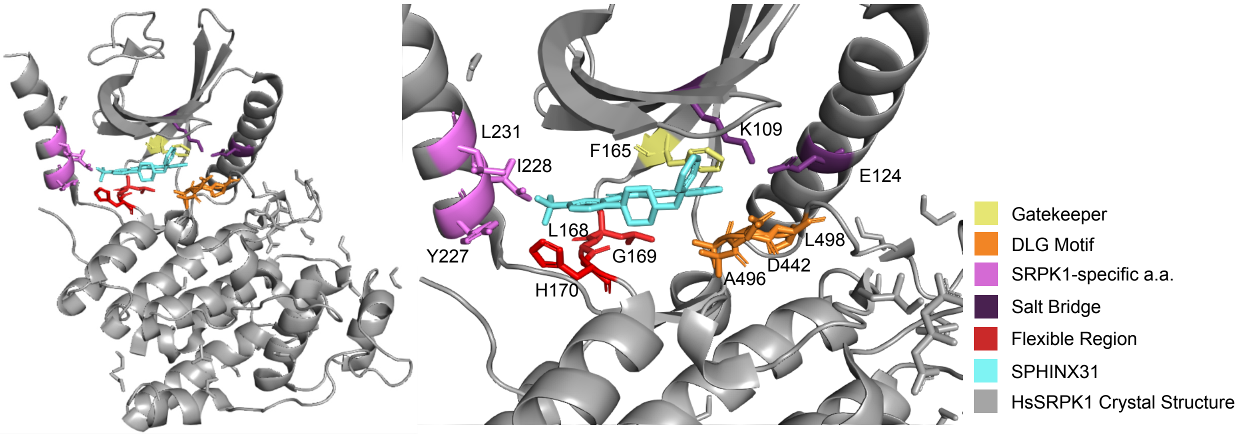


***Supplemental Figure 3. Crystal structure of HsSRPK1 with SPHINX31 in the active site.*** *Amino acids interacting with SPHINX31 are highlighted into groups and were acquired from Batson et al., 2017. Right panel shows a magnified view of the active site region.*

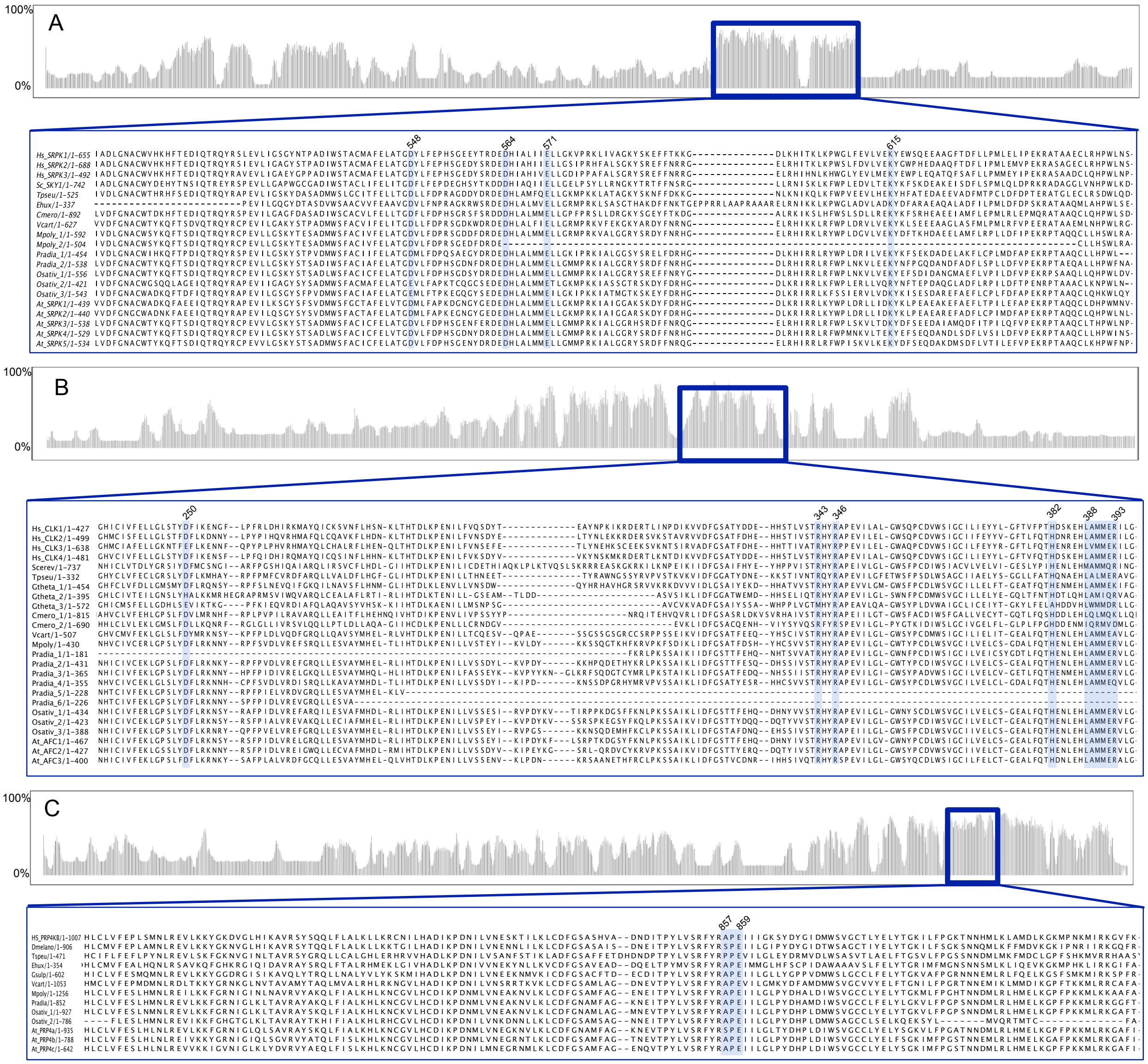


**Supplemental Figure 4. Protein alignment and percent conservation of human splicing-related protein kinases to orthologs from various species.** *Representative organisms from various taxa were selected for alignment. Alignment was performed using MAFFT with standard settings. Jalview software version 2.11.4 (https://www.jalview.org) was used to visualize alignment with grey bars representing percent conservation across the entire peptide sequence. Alignment area enclosed in a blue square represents region of the peptide sequence containing key amino acids required for kinase function. Highlighted blue columns represents amino acids required for kinase function. (A) SRPK alignment, (B) AFC alignment, and (C) PRP4Ks alignment.*

| | | | | | | |
|---|---|---|---|---|---|---|
| **AFC1** | 100% | | | | | |
| **AFC2** | 71.42% | 100% | | | | |
| **AFC3** | 65% | 60.75% | 100% | | | |
| **CLK1** | 35.54% | 40.74% | 40.75% | 100% | | |
| **CLK22** | 37.47% | 43.79% | 42.5% | 54.75% | 100% | |
| **CLK3** | 34.68% | 38.17% | 40.25% | 48.76% | 61.22% | 100% |
| | **AFC1** | **AFC2** | **AFC3** | **CLK1** | **CLK22** | **CLK3** |

***Supplemental Figure 5. AtAFC protein sequence percent identity comparisons to HsCLKs.*** *peptide sequences of AtAFCs and HsCLKs were aligned using MAFFT with default settings. Resulting alignment was inputted through the Sequence Identities and Similarities (SIAS) tool (http://imed.med.ucm.es/Tools/sias.html) with BLOSUM 62 as the scoring matrix.*

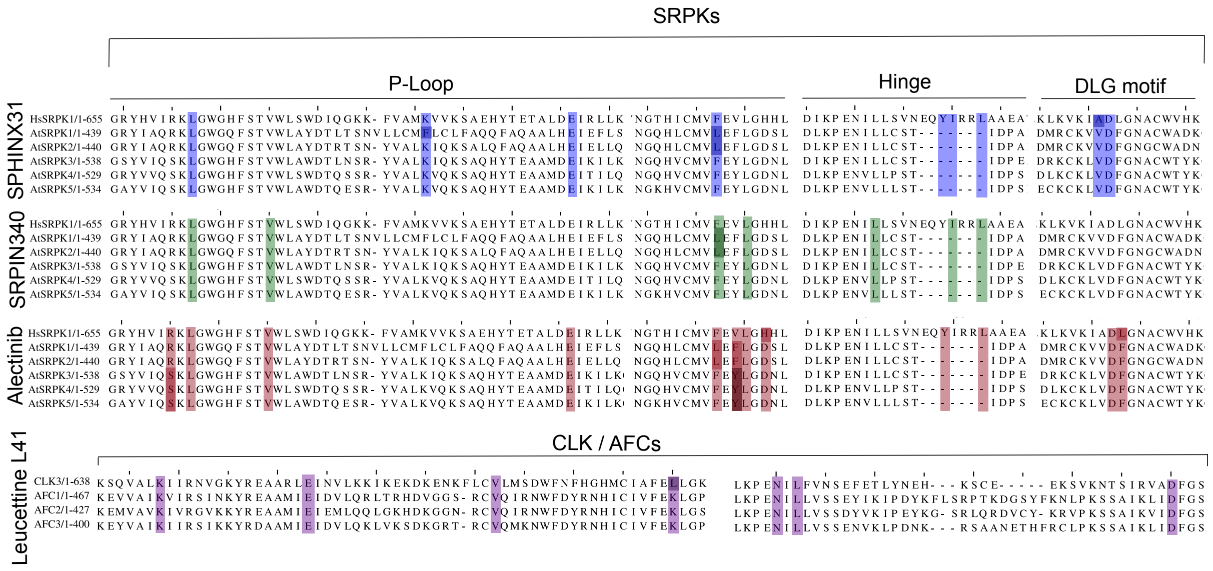


***Supplemental Figure 6. Protein sequence alignment of HsSRPK to AtSRPK depicting conservation amongst the amino acid targeted by each of the four chemical inhibitors.*** *Highlighted columns represent amino acids that were reported in literature (SPHINX31; Baston et al. 2017, SRPIN340; Marooka et al., 2015, Leucettine L41; Debdab et al., 2011, Alectinib; Hatcher et al., 2018) to be targeted by each chemical inhibitor. Peptide sequences were aligned with MAFFT with standard settings. Specific amino acid targeted by Leucettine L41 have only been investigated in HsCLKs despite showing inhibitory effects on other splicing-related protein kinases, thus only AFCs amino acid conversation is shown.*

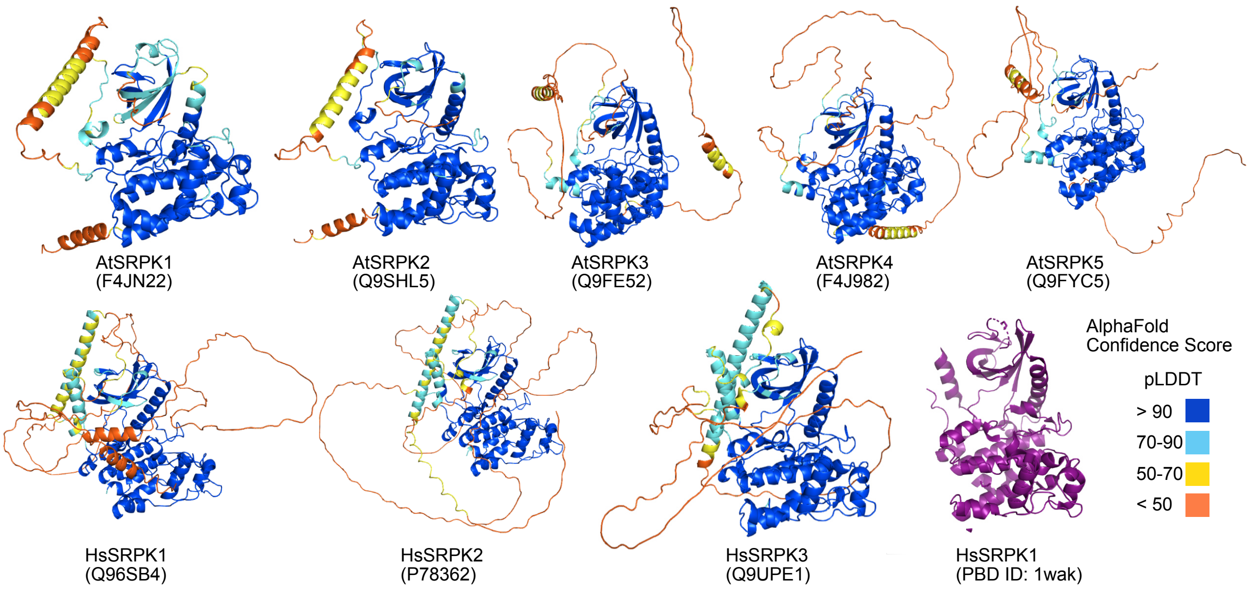


***Supplemental Figure 7. AtSRPK and HsSRPK 3-dimensional structure visualized with Alphafold confidence scores.*** *AtSRPK 3D structure were acquired from Alphafold (https://alphafold.ebi.ac.uk) and visualized using Pymol vers 3.1 (https://www.pymol.org).*

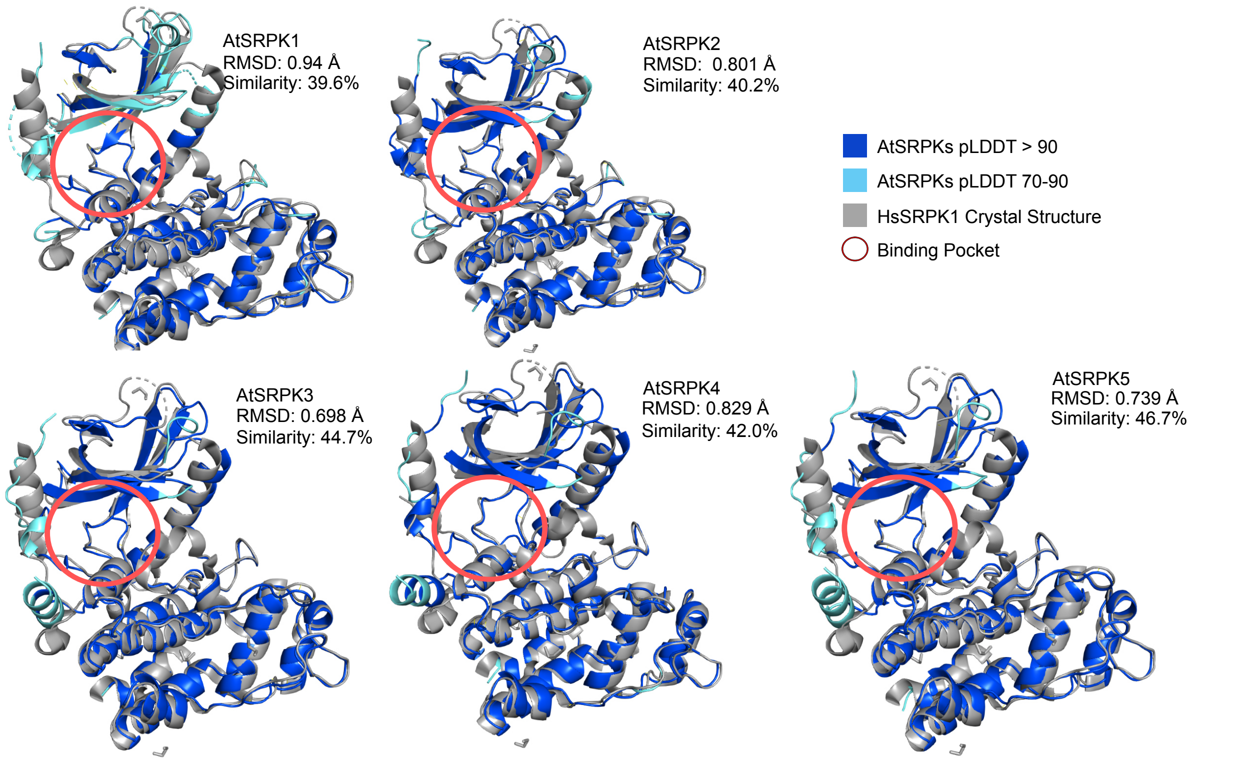


***Supplemental Figure 8. AtSRPK family superimposed on HsSRPK1 highlighting structural confirmation**. Structural portions below 70 pLDDT confidence scores were removed and then aligned to HsSRPK1 in PyMol vers 3.1 (https://www.pymol.org). Grey represent HsSRPK1 crystal structure. Blue represents AtSRPK with pLDDT > 90 and light blue is AtSRPK with pLDDT score between 70-90. Red circles highlights active site.*

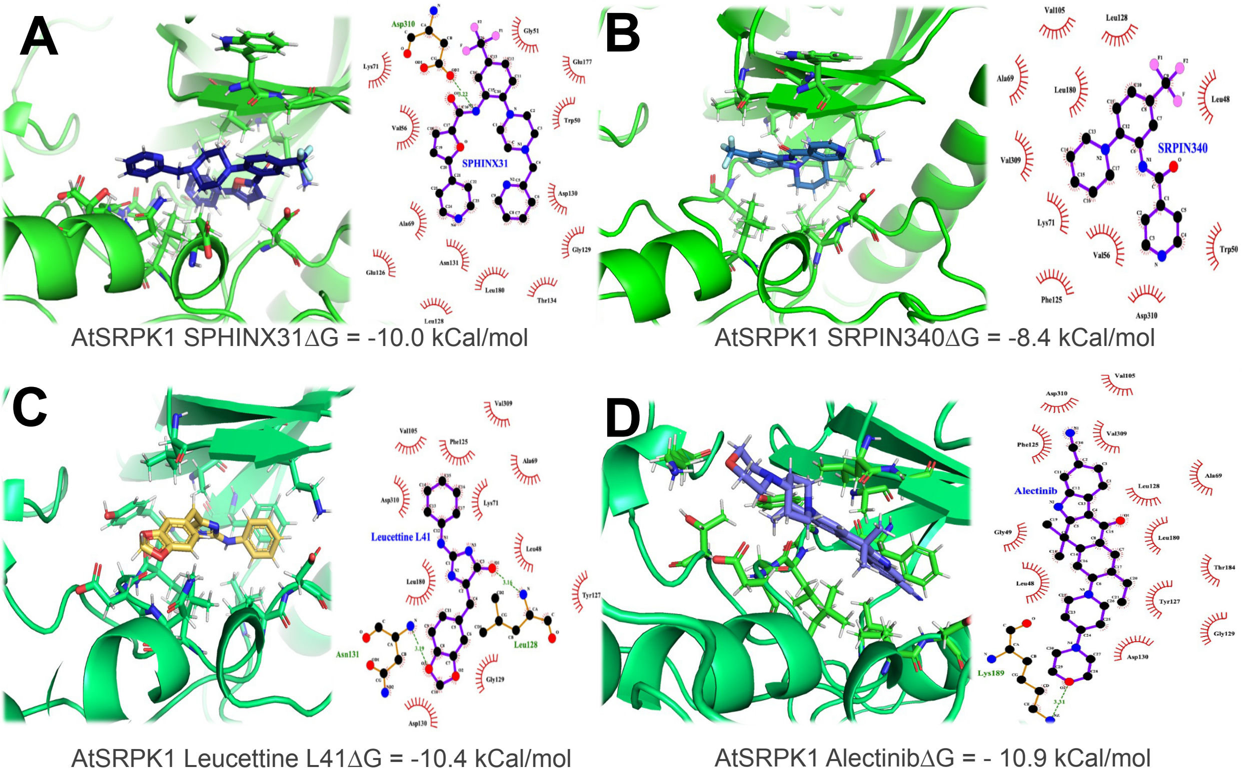


***Supplemental Figure 9. AtSRPK1 predicted interaction with the four chemical inhibitors.*** *AtSRPK1 predicted structure was imported from Alphafold and inhibitor's chemical structure was acquired from crystal structure PDB files (SPHINX31: 5my8, SPRIN340: 4wua, Leucettine L41: 8P05, Alectinib:5XV7). Amino acids predicted to interact with chemical inhibitors were visualized using LigPlot (https://www.ebi.ac.uk/thornton-srv/software/LigPlus/).*

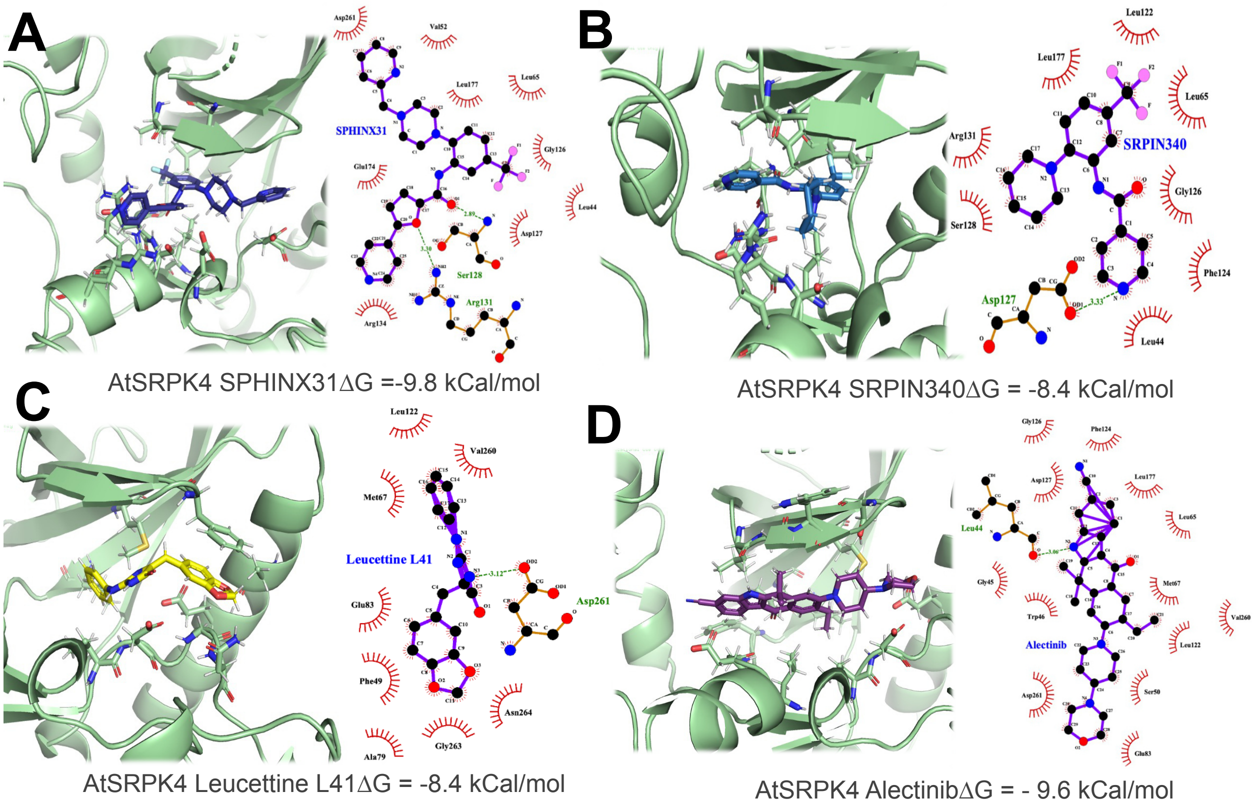


***Supplemental Figure 10. AtSRPK4 predicted interaction with the four chemical inhibitors.*** *AtSRPK4 predicted structure was imported from Alphafold and inhibitor's chemical structure was acquired from crystal structure PDB files (SPHINX31: 5my8, SPRIN340: 4wua, Leucettine L41: 8P05, Alectinib:5XV7). Amino acids predicted to interact with chemical inhibitors were visualized using LigPlot ([https://www.ebi.ac.uk/thornton-srv/software/LigPlus/](https://www.ebi.ac.uk/thornton-srv/software/LigPlus/)).*

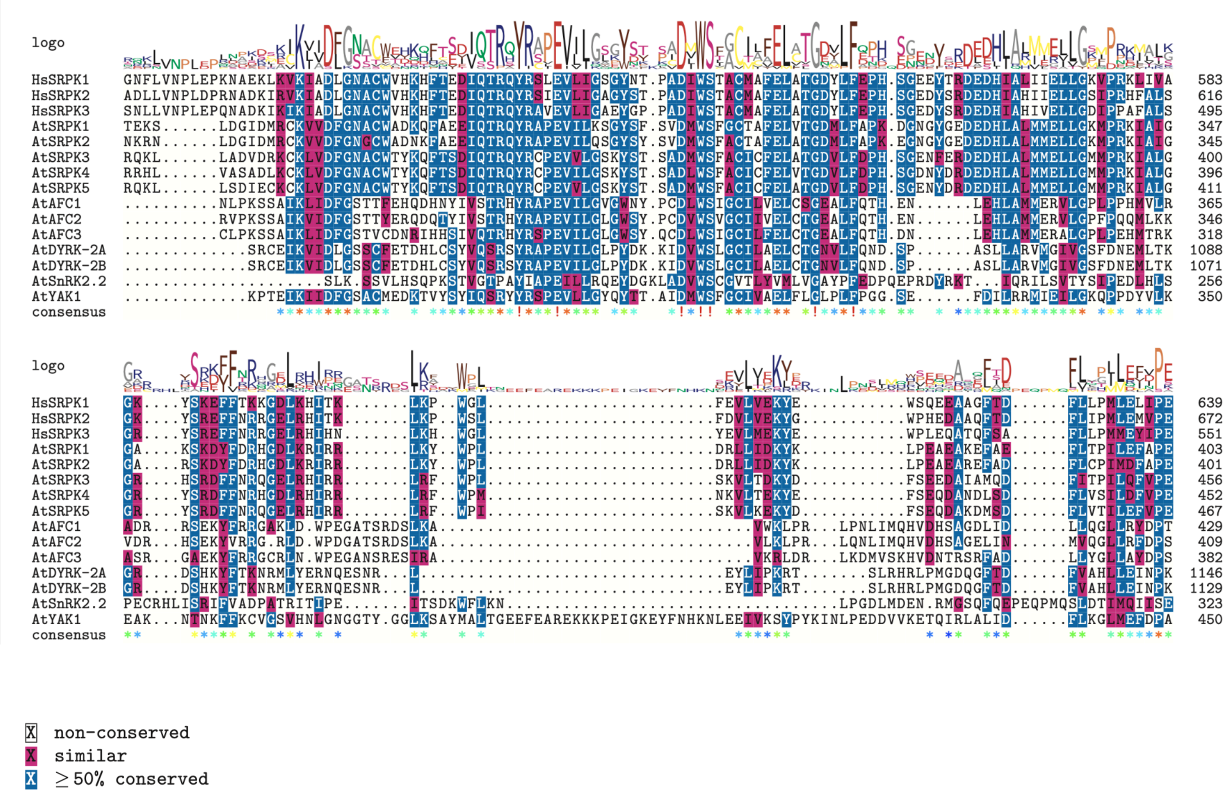


***Supplemental Figure 11. Multiple protein sequence alignment of HsSRPKs and Arabidopsis splicing-related kinases (AtSRPKs and AtAFCs), alongside other Arabidopsis proteins.*** *HsSRPK1 peptide sequence was BLASTp searched against the Arabidopsis proteome. Top hits that did not include AtSRPKs and AtAFCs were selected for MSA (AtDYRK and AtYAK1). Pink residues are similar while blue residues are greater than 50% conserved across all proteins.*

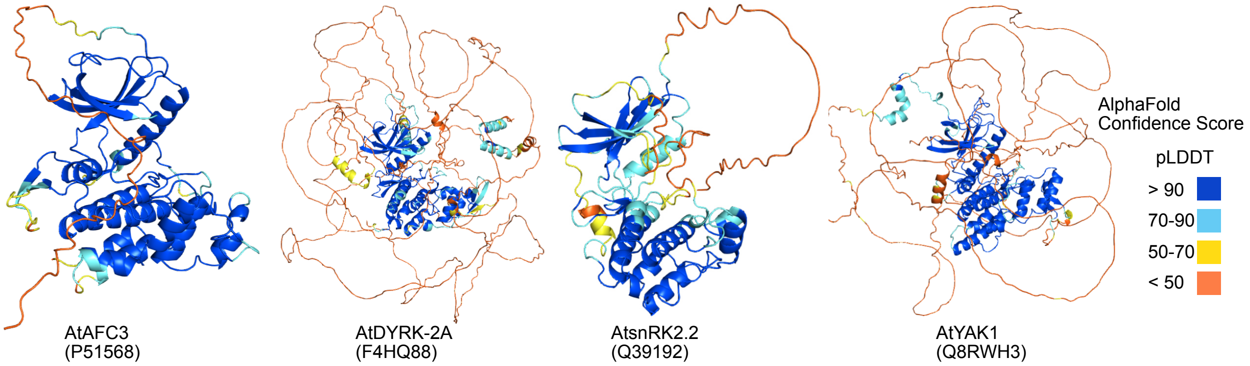


***Supplemental Figure 12. 3-dimensional structures of out-group Arabidopsis protein kinases visualized with Alphafold.*** *BLASTP analysis of the Arabidopsis proteome located proteins kinases with sequence similarity to SRPKs. Top scoring proteins were searched in Alphafold (https://alphafold.ebi.ac.uk) to acquire predicted 3D structure Alphafold and visualized using Pymol vers 3.1 (https://www.pymol.org).*

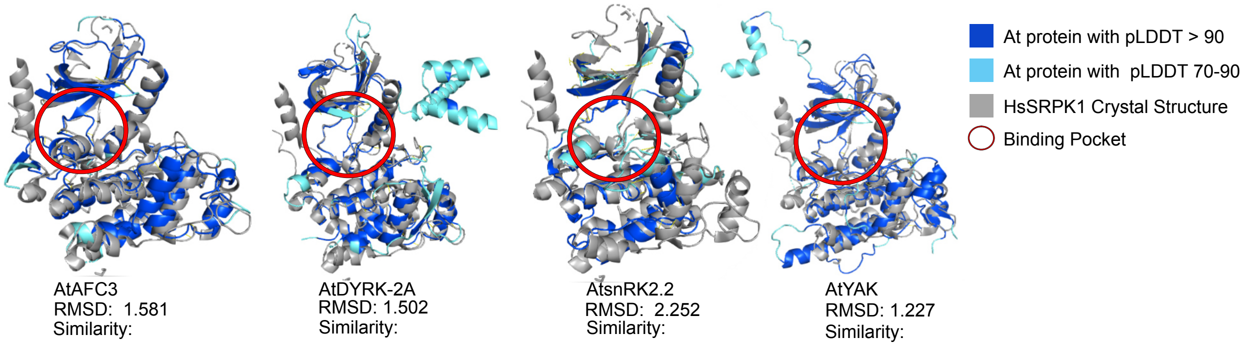


***Supplemental Figure 13. Out-group of Arabidopsis protein kinases superimposed on HsSRPK1 highlight structural confirmation conservation**. BLASTP of HsSRPK1 against the Arabidopsis proteome revealed proteins that have sequence similarity to HsSRPKs. Top scoring proteins were searched in Alphafold (https://alphafold.ebi.ac.uk) to acquire predicted 3D structure Alphafold and visualized using Pymol vers 3.1 (https://www.pymol.org). Structural portions below 70 pLDDT confidence scores were removed and then aligned to HsSRPK1. Blue represents AtSRPK with pLDDT > 90 and light blue is ATSRPK with pLDDT score between 70-90. Red circles highlights active site.*

Docking Simulation 1

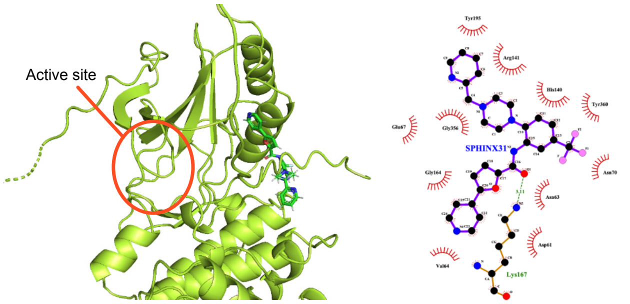


AtSnRK2.2 SPHINX31 ΔG = -8.8 kCal/mol

Docking Simulation 3

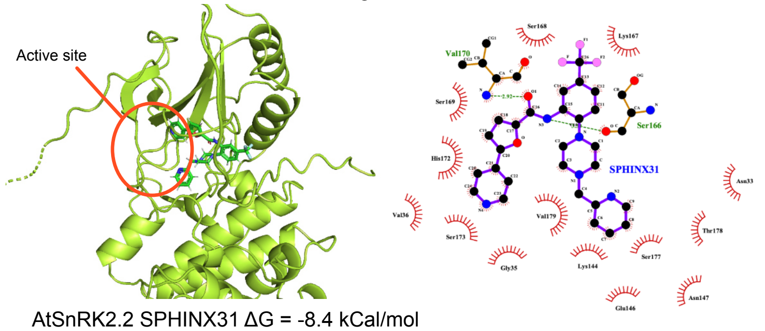


AtSnRK2.2 SPHINX31 ΔG = -8.4 kCal/mol

***Supplemental Figure 14. AtSnRK2.2 predicted interaction with SPHINX31.*** *AtSnRK2.2 predicted structure was imported from Alphafold and inhibitor's chemical structure was acquired from crystal structure PDB files (SPHINX31: 5my8). Amino acids predicted to interact with chemical inhibitors were visualized using LigPlot ([https://www.ebi.ac.uk/thornton-srv/software/LigPlus/](https://www.ebi.ac.uk/thornton-srv/software/LigPlus/)).*

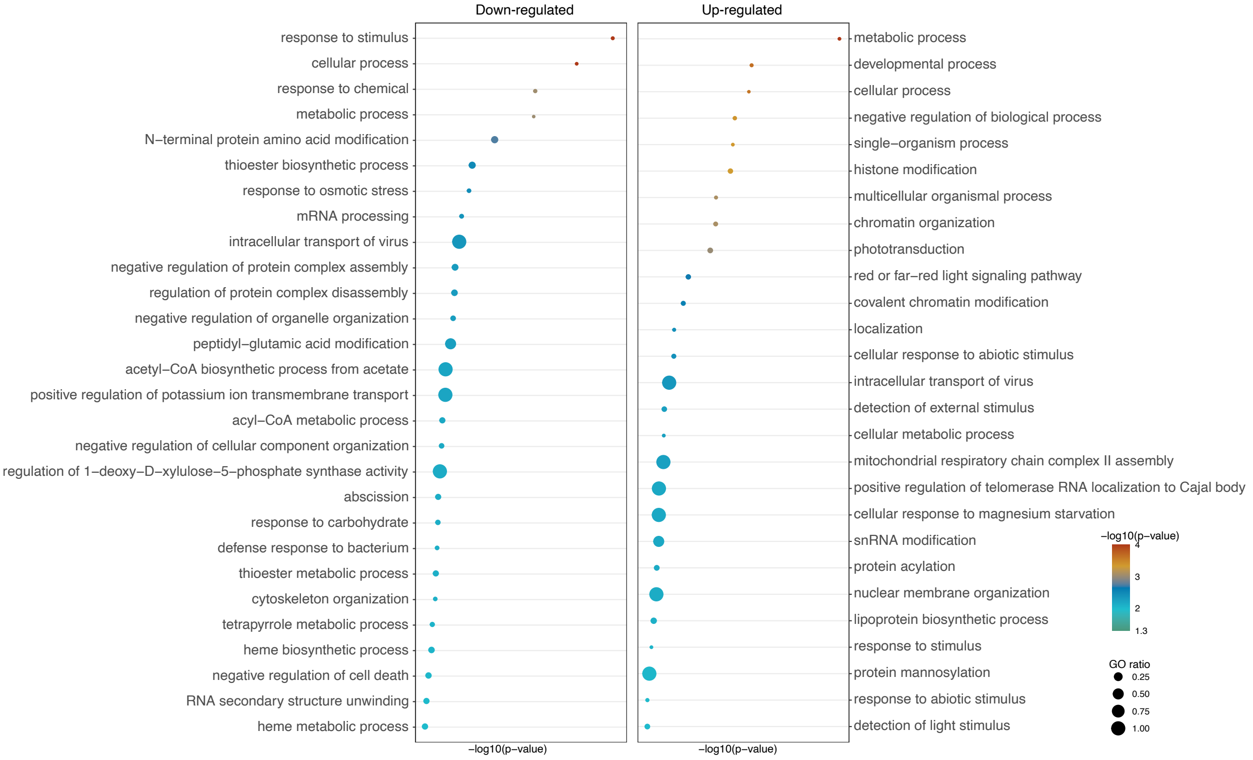


***Supplemental Figure 15. Biological gene ontology enrichment of A3SS alternative spliced genes in SPHINX31 inhibited Arabidopsis seedlings.*** *AS events were detected using ASTools with default settings (Qi et al., 2022; http://zzdlab.com/ASTool/). Up-regulated (ΔPSI > 0.1) and down-regulated (ΔPSI < 0.1) A3SS events were queried against whole transcriptome background for GO enrichment. Results were filtered to biological processes and with a p-value < 0.01. GO ratio was calculated by dividing study term by population term.*

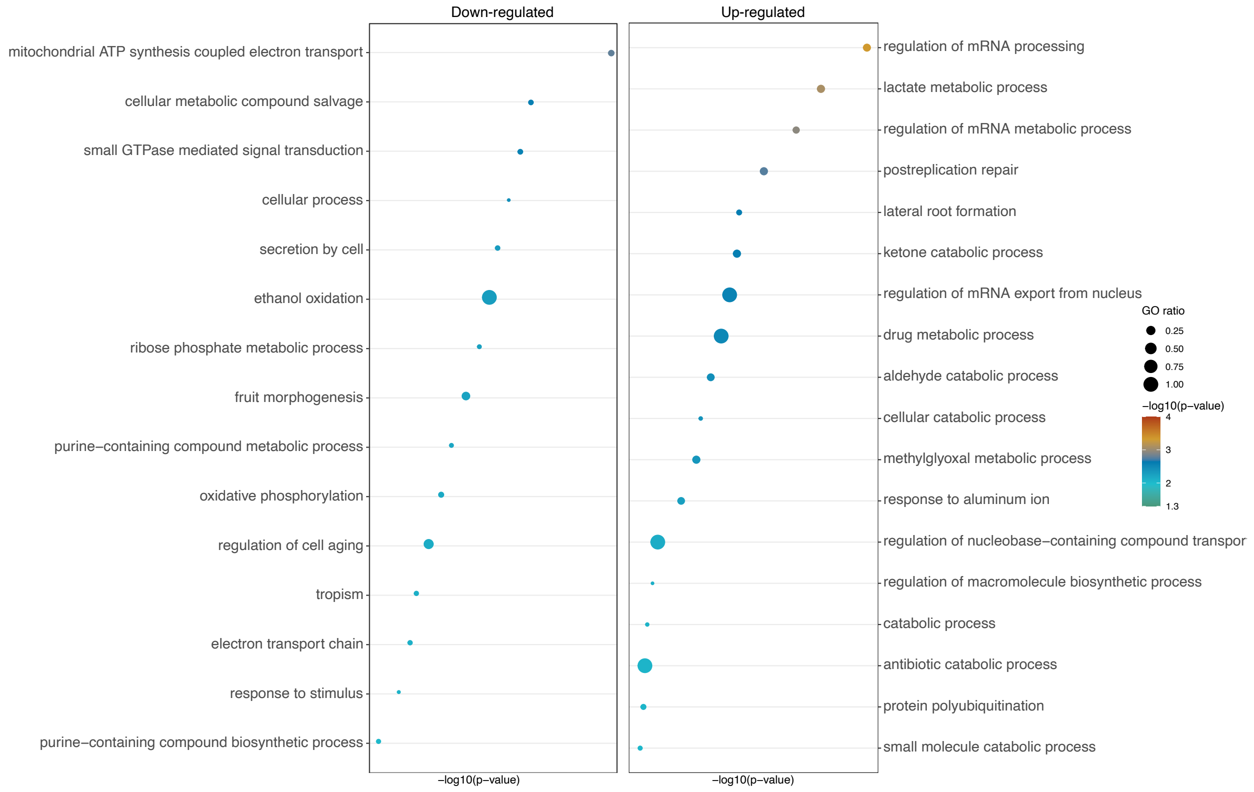


***Supplemental Figure 16. Gene ontology enrichment of A5SS alternative spliced genes in SPHINX31.*** *AS events were detected using ASTools with default settings (Qi et al., 2022; http://zzdlab.com/ASTool/). Up-regulated ˇ( PSI > 0.1) and down-regulated ˇ( PSI < 0.1) A5SS events were queried against whole transcriptome background for GO enrichment. Results were filtered to biological processes and with a p-value < 0.01. GO ratio was calculated by dividing study term by population term.*

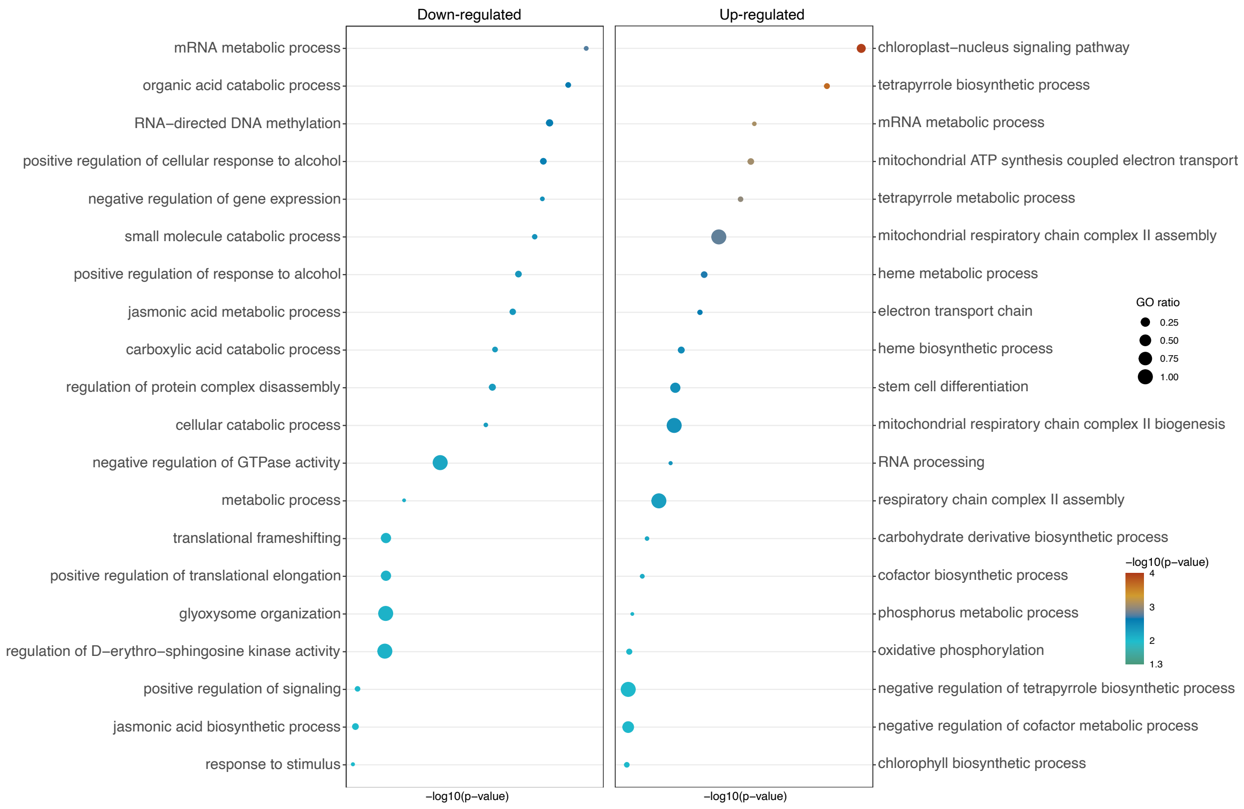


***Supplemental Figure 17. Gene ontology enrichment of ES alternative spliced genes in SPHINX31.*** *AS events were detected using ASTools with default settings (Qi et al., 2022; http://zzdlab.com/ASTool/). Up-regulated (ΔPSI > 0.1) and down-regulated (ΔPSI < 0.1) ES events were queried against whole transcriptome background for GO enrichment. Results were filtered to biological processes and with a p-value < 0.01. GO ratio was calculated by dividing study term by population term.*

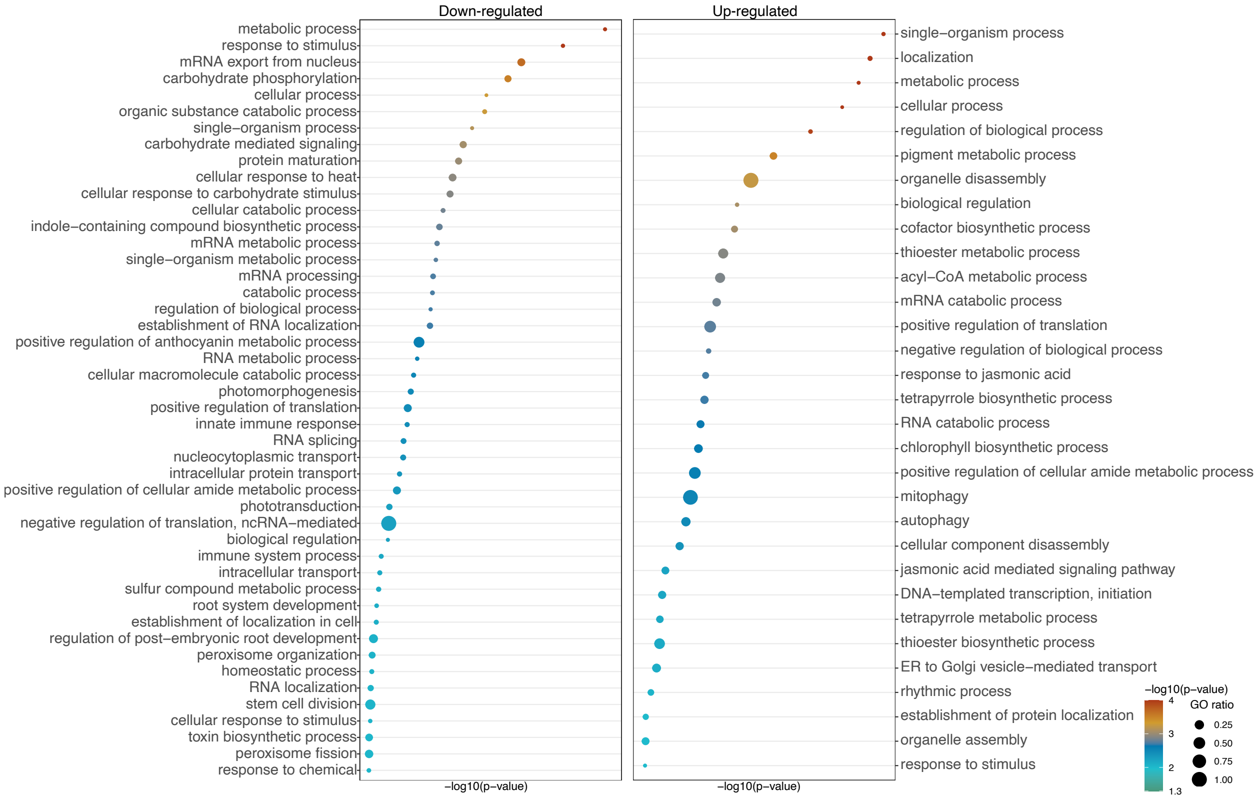


***Supplemental Figure 18. Gene ontology enrichment of IR alternative spliced genes in SPHINX31.*** *AS events were detected using ASTools with default settings (Qi et al., 2022; http://zzdlab.com/ASTool/). Up-regulated (ΔPSI > 0.1) and down-regulated (ΔPSI < 0.1) IR events were queried against whole transcriptome background for GO enrichment. Results were filtered to biological processes and with a p-value < 0.01. GO ratio was calculated by dividing study term by population term.*
